# DET1-mediated COP1 regulation avoids HY5 activity over second-site targets to tune plant photomorphogenesis

**DOI:** 10.1101/2020.09.30.318253

**Authors:** Esther Cañibano, Clara Bourbousse, Marta Garcia-Leon, Lea Wolff, Camila Garcia-Baudino, Fredy Barneche, Vicente Rubio, Sandra Fonseca

**Affiliations:** Centro Nacional de Biotecnología, CNB-CSIC, Madrid, 28049, Spain; Institut de biologie de l’Ecole normale supérieure (IBENS), Ecole normale supérieure, CNRS, INSERM, Université PSL, Paris, 75005, France

**Keywords:** HY5, DET1, COP1, photomorphogenesis, light signalling, *fusca*, de-etiolation, CRL4, C3D complex, CSN, PIF3, E3 ubiquitin ligase, ubiquitination, signalosome, proteasome, Arabidopsis

## Abstract

DE-ETIOLATED1 (DET1) is a negative regulator of plant photomorphogenesis acting as a component of the C3D complex, which can further associate to CULLIN4 to form a CRL4^C3D^ E3 ubiquitin ligase. CRL4^C3D^ is thought to act together with CRL4^COP1SPA^ ubiquitin ligase, to promote the ubiquitin-mediated degradation of the master regulatory transcription factor ELONGATED HYPOCOTYL5 (HY5), thereby controlling photomorphogenic gene regulatory networks. Yet, functional links between COP1 and DET1 have long remained elusive. Here, upon mass spectrometry identification of DET1 and COP1-associated proteins, we provide *in vivo* evidence that DET1 associates with COP1 to promote its destabilization, a process necessary to dampen HY5 protein abundance. By regulating HY5 over-accumulation, DET1 is critical to avoid its association to second-site loci, including many PIF3 target genes. Accordingly, excessive HY5 levels result in an increased HY5 repressive activity and are sufficient to trigger *fusca*-like phenotypes otherwise observed typically in *COP1* and *COP9* signalosome mutant seedlings. This study therefore identifies that DET1-mediated regulation of COP1 stability tunes down HY5 cistrome and avoids hyper-photomorphogenic responses that might compromise plant viability.

## Introduction

Light fuels plant life and is an essential cue that modulates growth and development throughout all the plant life cycle. Initial exposure of a young or germinating seedling to light sensed by photoreceptors initiates a light signalling cascade that triggers the passage from skotomorphogenic to photomorphogenic growth (Von Arnim and Deng, 1996).

To identify genes controlling plant photomorphogenesis, independent screenings were performed in the 90’s, which led to the isolation of the *det* (deetiolated) and *cop* (constitutive photomorphogenic) mutant families (Chory *et al*., 1989; Deng *et al*., 1991). Many of these mutants are allelic to the *fusca* (*fus*) mutants that excessively accumulate anthocyanin in embryos and seedlings in both light and dark (Castle and Meinke, 1994; Misera *et al*., 1994; Pepper *et al*., 1994). COP/DET/FUS group of proteins contain components of CULLIN4 based E3 ubiquitin ligases (CRL4) that mediate protein polyubiquitination marking for proteasomal degradation, and members of the signalosome (CSN). CSN is a multimeric complex (CSN1-8) structurally similar to the proteasome lid that enables deconjugation of NEDD8/RUB1 from CRLs, a process essential for CRL inactivation and recycling (Wei *et al*., 1994; Lyapina *et al*., 2001; Serino and Deng, 2003; Lau and Deng, 2012; Qin *et al*., 2020).

In Arabidopsis, COP1 is a central repressor of photomorphogenesis that works in complex with SUPPRESSOR OF PHYA-105 (SPA1-SPA4) proteins (Laubinger and Hoecker, 2003; Saijo *et al*., 2003; Seo *et al*., 2003; Laubinger *et al*., 2004). Light exposure reduces COP1 activity by controlling its transcript accumulation (Zhu *et al*., 2008; Huang *et al*., 2012); its exclusion from the nucleus (von Arnim and Deng, 1994; von Arnim *et al*., 1997; Pacín *et al*., 2014); degradation of its partner SPA2 (Balcerowicz *et al*., 2011; Chen *et al*., 2015) and by promoting its inactivation through association with photoreceptors (Huang *et al*., 2014; Podolec and Ulm, 2018). Though COP1 itself displays E3 ubiquitin ligase activity in vitro, which might promote its degradation according to results obtained for human COP1, it further associates *in vivo* with CUL4 to form CRL4^COP1SPA^ E3 ubiquitin ligase complexes (Osterlund *et al*., 2000; Saijo *et al*., 2003; Seo *et al*., 2003; Dornan *et al*., 2006; Chen *et al*., 2010). COP1 is highly active in darkness and targets numerous transcription factors (TFs) for proteasomal degradation, many of them being positive regulators of photomorphogenesis, such as ELONGATED HYPOCOTYL5 (HY5) and its homolog HYH, HIGH IN FAR RED 1 (HFR1) and LONG AFTER FAR-RED LIGHT 1 (LAF1) (Hardtke and Deng, 2000; Osterlund *et al*., 2000; Holm *et al*., 2002; Seo *et al*., 2003; Jang *et al*., 2005; Yang *et al*., 2005; Hoecker, 2017). Hence, a key function of COP1 is to regulate the availability of photo/skoto-morphogenic TFs including HY5.

HY5 is a basic leucine zipper (bZIP) TF that promotes photomorphogenesis by regulating, directly or indirectly, the expression of as much as one-third of Arabidopsis genes involved in diverse hormonal and metabolic pathways (Koornneef *et al*., 1980; Oyama *et al*., 1997; Jakoby *et al*., 2002; Lee *et al*., 2007; Zhang *et al*., 2011). In darkness, HY5 is a substrate of CRL4^COP1SPA^, and upon COP1 inactivation by light, HY5 accumulates in a proportional manner with light intensity (Osterlund *et al*., 2000). HY5 also positively regulates its own gene expression and directly binds to the promoters of anthocyanin, carotenoid and chlorophyll biosynthetic genes (Abbas *et al*., 2014; Binkert *et al*., 2014; Gangappa and Botto, 2016). Although HY5 does not bear an activation or repression domain (Ang *et al*., 1998), it mainly behaves *in vivo* as a transcriptional activator (Burko *et al*., 2020). Like other TFs regulating photomorphogenesis such as the PIFs, HY5 selectively binds to G-box sequence motifs (CACGTG) and variants (Gangappa and Botto, 2016). Attempts to determine the genes directly targeted by HY5 using different approaches (ChIP-chip, ChIP-seq), antibodies and transgene-driven expression levels gave rise to highly variable results ranging from 297 to 11797 genes (Lee *et al*., 2007; Zhang *et al*., 2011; Kurihara *et al*., 2014; Hajdu *et al*., 2018; Burko *et al*., 2020). HY5 acts in concert with other TFs to activate specific targets in specific tissues and conditions (Shin *et al*., 2007; Gangappa and Botto, 2016), while in other cases a competition for binding to promoter sequences was described (Toledo-Ortiz *et al*., 2014; Xu *et al*., 2014; Li and He, 2016; Gangappa and Kumar, 2017; Nawkar *et al*., 2017). Hence, several evidences indicate that regulation of HY5 stability is important, as rate-limited or excess amounts of HY5 may differently influence its regulatory activity.

HY5 down-regulation has also been shown to rely on DE-ETIOLATED1 (DET1), a chromatin-associated protein that is also key for photomorphogenesis repression in darkness (Chory *et al*., 1989; Chory and Peto, 1990; Pepper *et al*., 1994; Benvenuto *et al*., 2002). Both *cop1* and *det1* mutant alleles accumulate high protein levels of HY5, supporting the idea that DET1 is necessary for COP1 function on HY5 degradation (Osterlund *et al*., 2000). Accordingly, *hy5* mutations partially suppress *det1* phenotypes (Pepper and Chory, 1997) but, still, the possibility of direct relationship between DET1 and HY5 remain elusive.

Together with DAMAGED DNA BINDING PROTEIN1 (<u>D</u>DB1), <u>C</u>OP10, and DDB1-ASSOCIATED1 (<u>D</u>DA1), <u>D</u>ET1 forms a C3D adaptor module (also termed CDDD) of CRL4^<u>C3D</u>^ E3 ubiquitin ligases to potentially target proteins for proteasomal degradation, including DDB2 (damaged DNA sensor), two subunits of a histone H2B deubiquitination module, the PYL8 abscisic acid receptor, and the HFR1 TF (Schroeder *et al*., 2002; Chen *et al*., 2006; Castells *et al*., 2011; Irigoyen *et al*., 2014; Nassrallah *et al*., 2018). Oppositely to its negative influence on HY5, DET1 contributes to the stabilization of PHYTOCHROME INTERACTIONG FACTORs (PIFs) TFs in the dark and can act as a transcriptional co-repressor (Maxwell *et al*., 2003; Lau *et al*., 2011; Dong *et al*., 2014; Shi *et al*., 2015). The repressive role of DET1 on transcription relies in part to its ability to regulate chromatin states. Plant DET1 shows high affinity for histone H2B and controls H2B ubiquitination levels on most genes mainly through proteolytic degradation of the DUBm (Benvenuto *et al*., 2002; Nassrallah *et al*., 2018). More generally, DET1 and COP1 are evolutionarily conserved in animals and plants (Schwechheimer and Deng, 2000; Yi and Deng, 2005; Olma *et al*., 2009). In humans however, the tumor-suppressor COP1 serves as an adaptor protein for DET1 to form the CRL4^DET1COP1^ complex. The latter recruits TFs such as c-Jun, ETS1/2 and ETV1/4/5, and target them for proteasomal degradation (Wertz *et al*., 2004; Vitari *et al*., 2011; Marine, 2012) Arabidopsis COP10 and COP1 were found to associate together (Yanagawa *et al*., 2004; Chen *et al*., 2010), but their functional relationship is plant systems remain elusive. Lack of evidence of physical association between DET1 and COP1 proteins conducted to the idea that they are part of distinct CRL4 complexes that cooperate in plant photomorphogenesis through an unidentified mechanism (Lau and Deng, 2012; Huang *et al*., 2014).

Here, we report that DET1 and COP1 can associate in the same complexes *in vivo*, DET1 being required for decreasing COP1 level in a light-independent manner. Despite this observation being seemingly paradoxical with HY5 over-accumulation in *det1-1* mutant seedlings, our observations suggest that DET1-mediated COP1 destabilization is necessary for HY5 degradation. By profiling HY5 chromatin landscape in wild-type and in *det1-1* and transgenic lines that over-accumulate HY5 protein, we further showed that uncontrolled HY5 levels lead to an aberrant enrichment over the promoter of second-site gene targets shared by light-regulated TFs such as PIF3, and frequently induce *fusca*-like phenotypes. Collectively, these observations led us to propose a direct role for DET1 in the regulation of COP1-mediated HY5 protein regulation to tune its association to a potentially very large gene repertoire. This study further identifies dynamic interaction of HY5 with hundreds’ genomic loci that may underlie its interplay with other TFs such as PIF3.

## Results

### COP1 and DET1 co-exist in one or more protein complexes

In search for DET1 interactors, we carried out Tandem Affinity Purification (TAP) coupled to mass spectrometry (MS) analysis of DET1 constitutively expressed in Arabidopsis cell cultures grown under dark conditions. In total, five independent TAP assays in which cells were collected in dark or after a 24 h white light treatment, allowed identifying proteins that co-purified with DET1. In all experiments, a large number of peptides representative of all known components of the C3D complex (DET1, DDB1, DDA1, COP10) was detected (Fig. 1A, Table S1 and S2), suggesting that the C3D complex is stable under these conditions as previously reported (Schroeder *et al*., 2002; Yanagawa *et al*., 2004; Olma *et al*., 2009; Irigoyen *et al*., 2014). Remarkably, the scores obtained for DDB1 (either DDB1a or DDB1b) recovery are in some experiments above those obtained for DET1, indicating that the large majority of DET1 proteins exist in association with DDB1. We also detected CUL4 peptides in high abundance, as well as all CSN subunits (CSN1 to 8) (Fig. 1A, Table S1) confirming that the C3D complex forms a CUL4 based E3 ligase (CRL4^C3D^) and that the C3D complex works in close association with the signalosome similarly to mammalian systems (Wertz *et al*., 2004; Chen *et al*., 2006; Lau and Deng, 2012). Among CSN subunits, CSN1 displays higher protein scores that might be due to its larger molecular size (easier to detect by MS) or because it represents a conserved direct contact point with DDB1 as previously reported in the structural analysis of human CSN-CRL4 complexes (Fig. 1A) (Cavadini *et al*., 2016).

**Figure 1.**
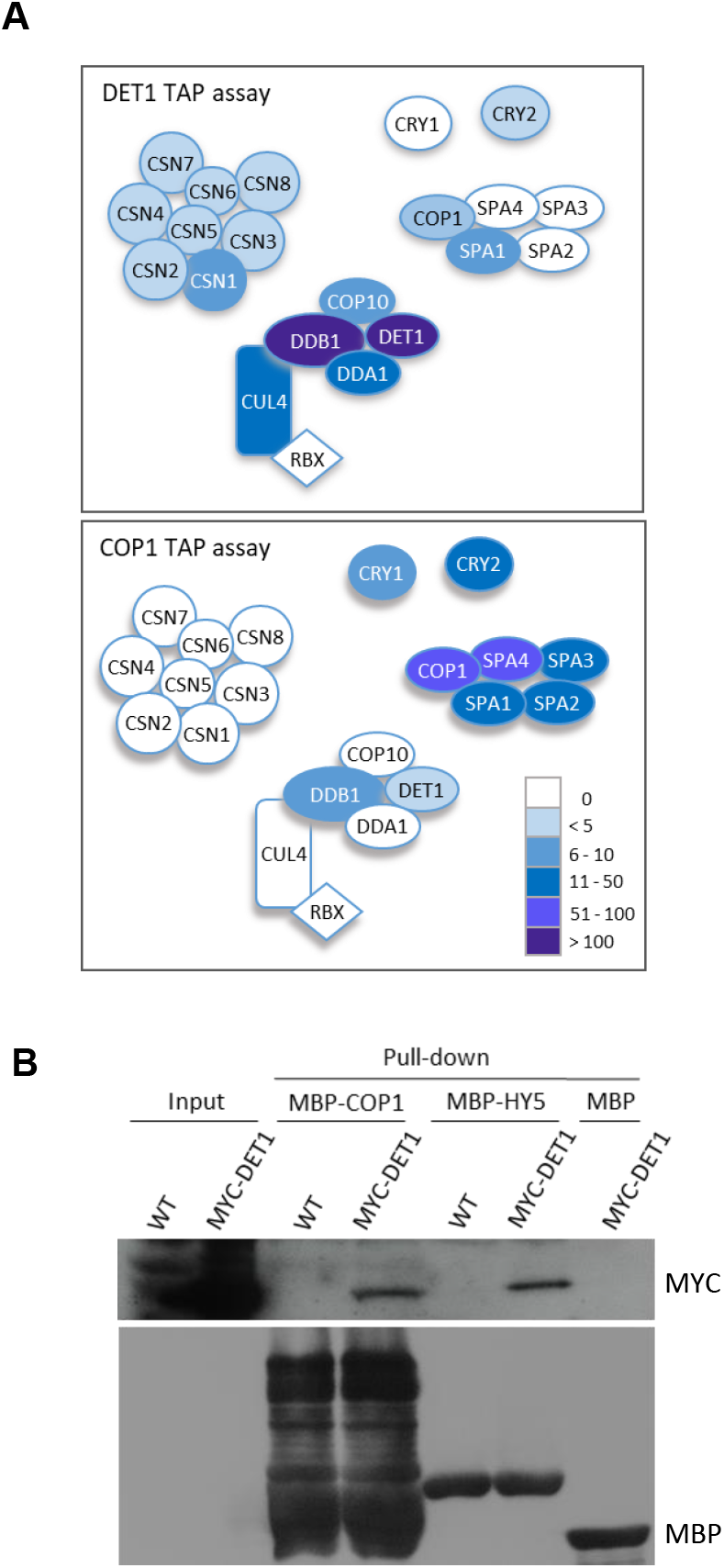
DET1 and COP1 associated proteins. (A) Schematic representation of proteins found to associate with DET1 and with COP1 in TAP assays. Colour code represents the maximum number of peptides for each represented protein found in a TAP assay as detailed in Table S1. DET1 and COP1 proteins were expressed in Arabidopsis cell cultures. Five independent TAP experiments were performed for DET1 and two for COP1 (Table S2). (B) MBP-COP1 and MBP-HY5 recombinant proteins expressed in *E. coli* pulled-down MYC-DET1 from 7-day old Arabidopsis seedlings. MBP recombinant protein was used as a control. Anti-MYC and anti-MBP antibodies were used for the immunoblots.

Strikingly, we detected COP1 and SPA1 peptides in three of the five DET1 TAP experiments (Fig. 1A, Table S1 and S2). Since a COP1-DET1 interaction was discarded long ago in Arabidopsis (Chen *et al*., 2010), we aimed to confirm this result by performing TAP assays for COP1. Expectedly, MS analysis of proteins co-purified with COP1 under dark and light conditions allowed the detection of a high number of peptides for all four SPA proteins (SPA1 to 4), and also from CRY1 and CRY2, confirming that COP1 predominantly associates with these proteins (Fig. 1A; Table S1, S3; Wang *et al*., 2001a; Yang *et al*., 2001; Saijo *et al*., 2003; Laubinger *et al*., 2004). Importantly, we also recovered peptides, at a relatively lower number, for DET1 and DDB1, indicating association of COP1 with the C3D complex. However, no peptides from C3D subunits COP10 and DDA1 were found, potentially owing to their smaller molecular size. Neither CSN nor CUL4 peptides were detected (Fig. 1, Table S1), which is noteworthy considering the proposal that COP1 binds DDB1 to form a stable CRL4^COP1SPA^ E3 ligase (Chen *et al*., 2010). From our data, we cannot discard the existence of these complexes, however, COP1 preferentially associates with SPA and CRY proteins instead of forming a stable CRL4^DDB1-COP1^ in our Arabidopsis cell suspensions, meaning that COP1 association with CUL4 is potentially transitory or cell-type specific.

Altogether, our results indicate that COP1 and DET1 associate transiently *in vivo*. This association was confirmed by semi-*in vivo* pull-down assays where bacteria-purified MBP-COP1 and MBP-HY5, but not the MBP alone, could pull-down MYC-DET1 from Arabidopsis protein extracts (Fig. 1B). These analyses support the existence of an association between DET1 and the COP1-HY5 module.

### DET1 represses COP1 accumulation in a light independent manner

Considering this association and the established role of DET1 in ubiquitin-mediated protein degradation, we hypothesized that DET1 could affect the accumulation of COP1. To test this, we analysed the accumulation of endogenous COP1 in wild-type (WT) and *det1-1* dark-grown seedlings and 3 or 24 hours upon exposure to light, as well as in plants germinated directly under light. We detected very low COP1 levels in dark-grown plants and slightly higher COP1 levels in light-grown plants, suggesting that low levels of COP1 in darkness are sufficient for its activity, including HY5 targeting for degradation (Fig. 2A).

**Figure 2.**
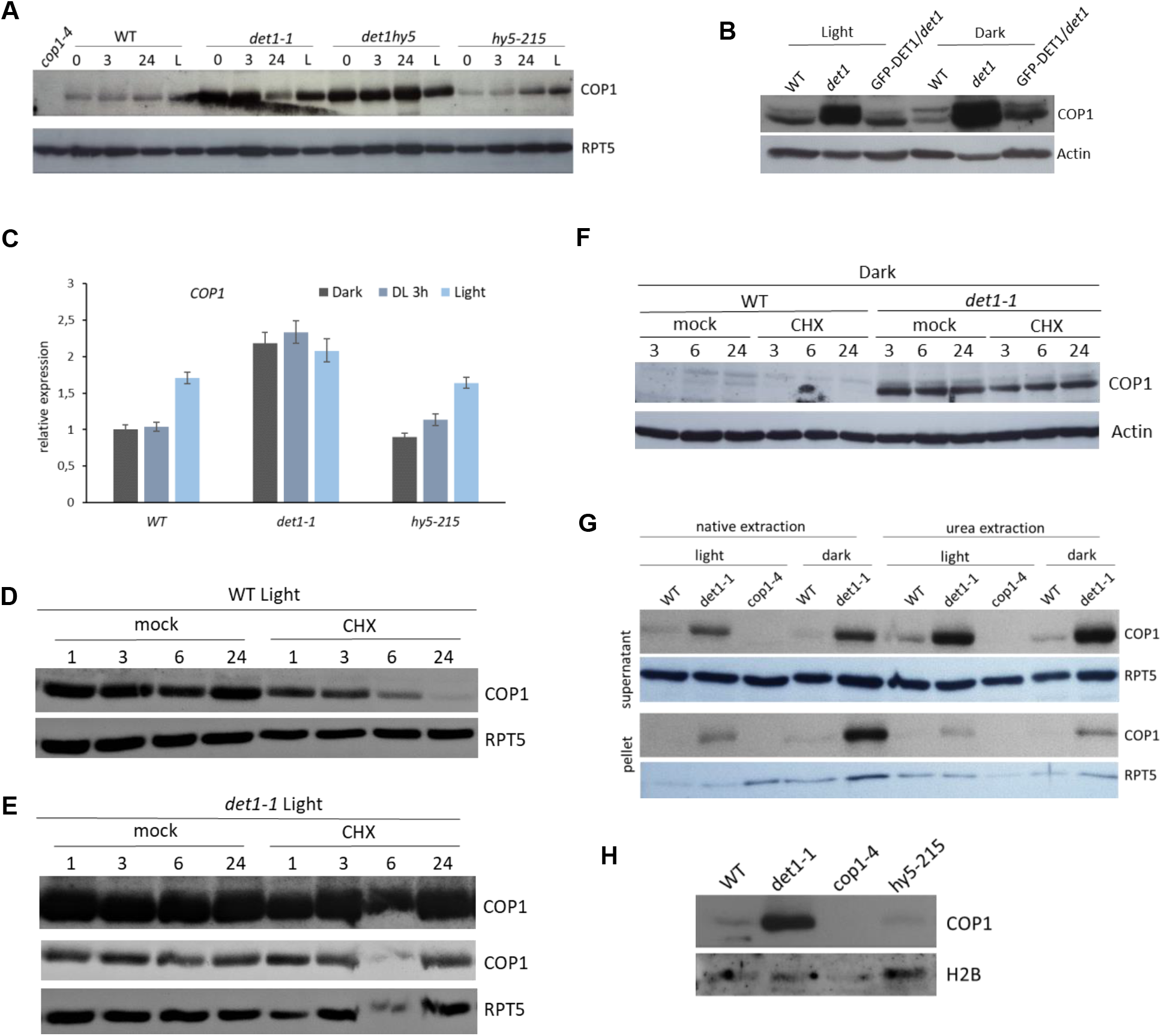
DET1 is necessary for COP1 destabilization. (A) Accumulation of endogenous COP1 (detected with anti-COP1) in WT *det1-1, det1-1 hy5-215* and *hy5-215* mutants grown in Dark (0) or under long day conditions for 6 days (L). Dark grown plants were transferred to light and collected after 3 or 24 hours. For loading control anti-RPT5 was used. (B) COP1 protein levels in complemented *GFP-DET1/det1-1* lines under light or dark conditions. (C) Analysis of *COP1* transcript levels relative to the levels of *COP1* in etiolated WT seedlings that were set as 1. *PP2A* levels were used as controls. Bars represent average ± SD of three replicates. Similar results were obtained by analysing different pools of plants as biological replicates. (D and E) COP1 protein levels in WT (D) or *det1-1* (E) after treatment with cycloheximide (CHX) versus plants treated with the solvent (mock control) for the indicated time under continuous light conditions. In (E) Blots are exposed at different times: the upper one corresponds to the same immunoblot and long exposure time as in (D) whereas a lower exposure blot is shown in the lower panel. (F) Cycloheximide treatment as in (D-E) in dark conditions. (G) COP1 accumulation in WT, *det1-1* and *cop1-4* in proteins extracts obtained using native and denaturing (4M Urea) extraction conditions. Soluble and insoluble fractions were loaded. RPT5 protein levels were detected as loading control. (H) COP1 accumulation in nuclear extracts of WT and indicated mutant backgrounds grown under long day conditions. Total Histone 2B (H2B) levels were used as loading control.

Surprisingly, COP1 protein accumulation in *det1-1* was much higher than in WT plants independently of the light condition used (Fig. 2A). Abundance of COP1 in *GFP-DET1/det1-1* complemented lines (Pepper and Chory, 1997) was similar to that of WT plants in both dark and light conditions, confirming that elevated COP1 levels are linked to *DET1* loss-of-function (Fig. 2B). It has been recently proposed that HY5 activity can induce COP1 accumulation (Burko *et al*., 2020). Since *det1-1* mutants display high levels of HY5 (Osterlund *et al*., 2000), we analysed COP1 accumulation in *hy5* and *det1hy5* mutants to discard a possible direct effect of HY5 over-accumulation on COP1 abundance. We found no difference between *hy5* and WT or between *det1hy5* and *det1-1*, demonstrating that COP1 over-accumulation in *det1-1* does not rely on *HY5* (Fig. 2A). In line with this observation, RT-qPCR analysis of *hy5* mutant seedlings confirmed that HY5 does not affect *COP1* gene expression in dark or light conditions (Fig. 2C; Fig. S1). In these transcript analyses, we observed a slight, yet significant increase in *COP1* RNA level in *det1-1* mutants, suggesting a primary negative influence of DET1 on *COP1* transcription. To test whether DET1 also controls COP1 accumulation at the post-transcriptional level, we treated Arabidopsis plants with the translation inhibitor cycloheximide (CHX) (Fig. 2D-F). We found that COP1 level strongly decreases in WT (Fig. 2D) within 6 hours after treatment while it remained mostly unaffected in *det1-1* mutant plants (Fig. 2E). This indicated that DET1 plays also a role in controlling COP1 protein stability. Similar results were obtained using dark-grown seedlings (Fig. 2F). Thus, DET1-mediated COP1 destabilization seems to be light independent.

Further considering the possibility that our native protein extraction may favour the extraction of the cytoplasmic pool of COP1 and that *det1-1* might alter COP1 nuclear accumulation in the dark (Chamovitz *et al*., 1996; von Arnim *et al*., 1997; Wang *et al*., 2009), we performed a denaturing protein extraction by adding 4M Urea to the extraction buffer and, in both cases, we analysed the insoluble (pellet) and soluble (supernatant) fractions. COP1 over-accumulated in *det1-1* in all the conditions tested (dark and light) even when the majority of proteins were recovered from the insoluble fraction (Fig. 2G). Since DET1 is a nuclear protein playing a role in the regulation of chromatin (Nassrallah *et al*., 2018), we performed a nuclear extraction to analyse COP1 nuclear level. We found that COP1 nuclear pool is much higher in *det1-1* background and is unaffected in *hy5* seedlings (Fig. 2H). Collectively, these analyses unveil a role for DET1 in moderating COP1 abundance at multiple levels, including protein destabilization.

### COP1 destabilization depends on the proteasome, C3D and CSN

Further supporting the idea that COP1 is being degraded by the proteasome independently of light, we observed that treatment with the proteasome inhibitor bortezomib (Bor) results in a moderate increase in COP1 protein accumulation in WT seedlings both under light and dark conditions (Fig. 3A and B). This was not the case in *det1-1* seedlings, Bor treatment having little or no effect on COP1 accumulation, suggesting that DET1 is mediating COP1 proteasomal degradation (Fig. 3A and B).

**Figure 3.**
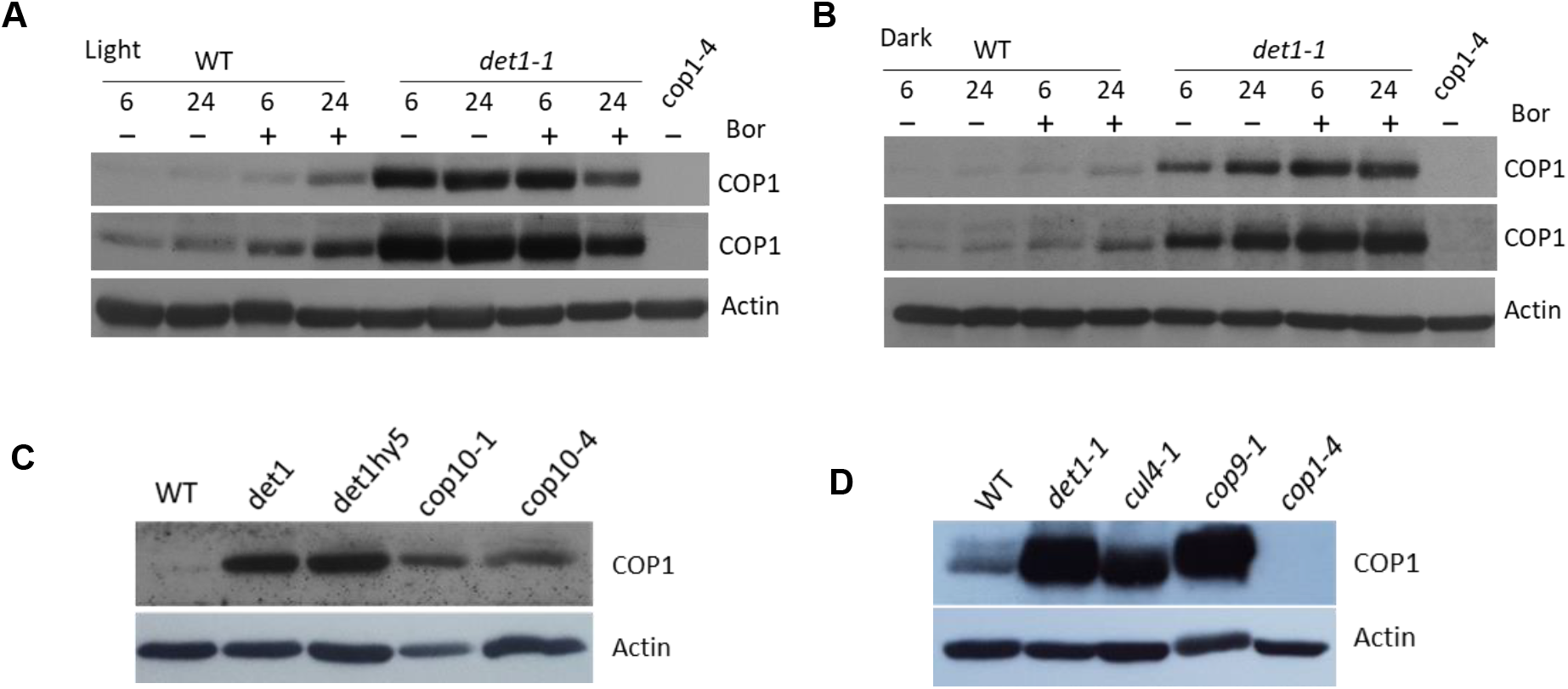
COP1 destabilization depends on an active proteasome and on functional CRL4^C3D^ and CSN complexes. (A and B) COP1 protein accumulation in WT and *det1-1* Arabidopsis seedlings treated with bortezomib (Bor) for 6 or 24 hours, under continuous light (A) or continuous dark conditions (B). (C) COP1 protein levels in different mutants grown under dark conditions. (D) COP1 protein levels in *cul4-1* mutant an in the homozygous *cop9-1 fusca* plants are similar to those on *det1-1*. All immunoblots were performed with anti-COP1 antibody and anti-actin or anti-RPT5 as loading controls.

DET1 and COP10 being part of the CUL4 based C3D complex, we further tested COP1 accumulation in *cop10* and *cul4* mutant seedlings. Both *cop10-4* (weak allele), *cop10-1* (null allele) and *cul4-1* seedlings displayed COP1 over-accumulation (Fig. 3C,D). As in TAP assays DET1 and the CSN associate together, we tested COP1 accumulation in *cop9-1* homozygous mutant for *CSN8* (Fig. 3D). In this mutant, COP1 accumulated to levels similar to those in *det1-1*, supporting the idea that CSN, through the canonical mechanism of CUL4 deneddylation, is required for COP1 rapid destabilization, likely by facilitating the function of a CRL4^C3D^.

### DET1 facilitates HY5 degradation

A role for DET1 in COP1 protein regulation represents a new hint on the early hypothesis that DET1 facilitates COP1 function in promoting HY5 degradation (Osterlund *et al*., 2000). Still, over-accumulation of HY5 in *det1-1* mutant plants having never been analysed in detail, we monitored HY5 accumulation in WT, *det1-1* and *cop1-4* seedlings for 24 h during de-etiolation. As reported earlier, in WT seedlings HY5 protein was detectable by immunoblot 3 h after illumination. HY5 also accumulates at high levels in dark-grown *det1-1* seedlings, and this accumulation is even increased after light exposure in a similar way than in *cop1* seedlings (Fig. 4A).

**Figure 4.**
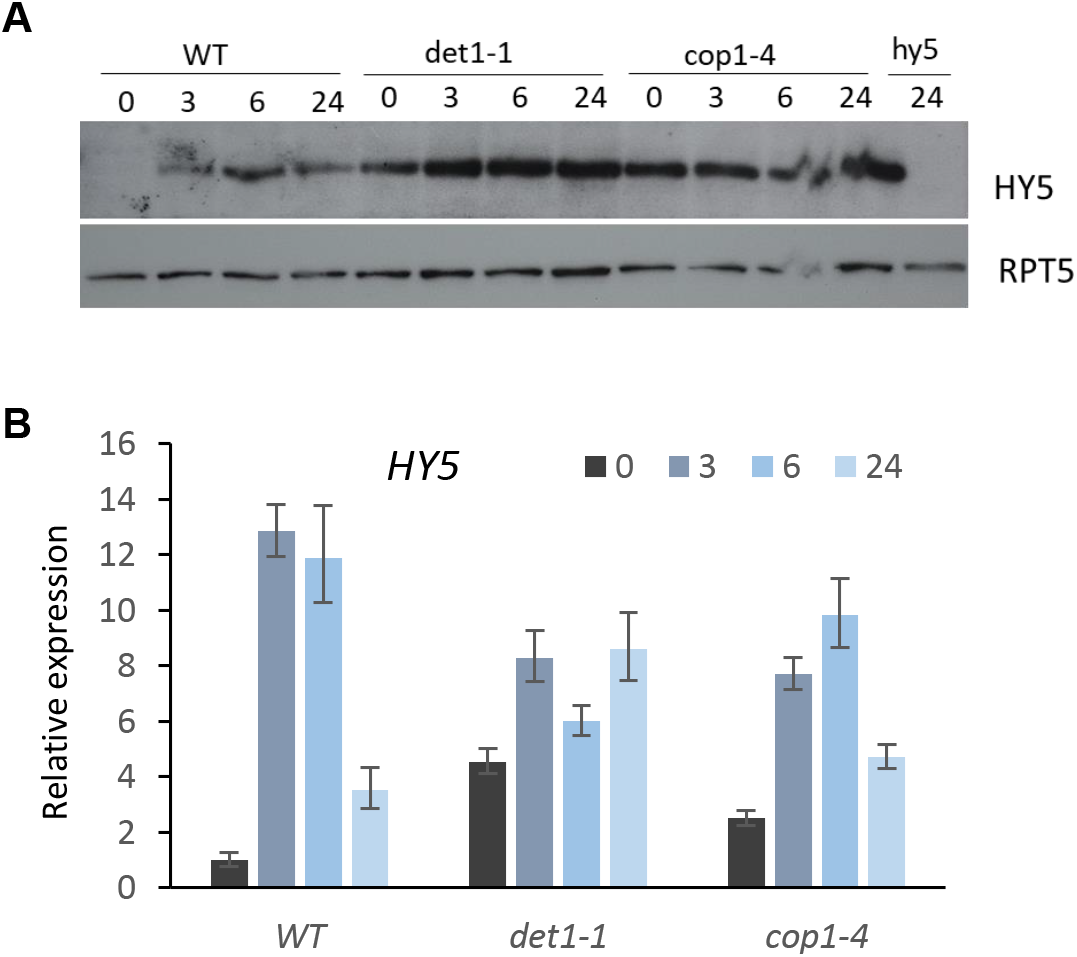
DET1 represses HY5 accumulation. (A) HY5 protein accumulation in Arabidopsis WT, *det1-1* and *cop1-4* seedlings germinated in light and, 3, 6 or 24 hours after transfer to light. Protein extracts from *hy5-215* mutant obtained 24 hours after transference to light was used as control. (B) Relative accumulation of *HY5* transcripts in Arabidopsis seedlings treated as in (A) analysed by qRT-PCR. The expression level of each gene was normalized to that of *ACTIN8* (*ACT8*). Expression levels for each gene are shown relative to the expression levels in WT under dark conditions, which is set as 1. Bars represent average ± SD from three replicates and the experiment was repeated with different pools of plants with similar results.

To test for a potential transcriptional regulation of *HY5* by DET1, we analysed *HY5* transcript levels in this experimental setup by qRT-PCR. As previously reported, *HY5* transcript level increased after exposure to light (Fig. 4B; Osterlund *et al*., 2000). In *det1-1* and *cop1-4* seedlings, *HY5* transcript accumulation shows little variations in response to light. Hence, based on these observations, as previously suggested (Osterlund *et al*., 2000), the discrepancy in the kinetics and low increase of *HY5* transcripts accumulation in these mutants does not appear to be on its own a major determinant of high HY5 protein over-accumulation in *det1-1* mutant plants.

Collectively, these findings shed light on intricate links between DET1 and COP1-HY5 protein abundance, suggesting that DET1 promotes COP1 destabilization and, yet, is further necessary for COP1 activity in HY5 degradation.

### HY5 over-accumulation increases its chromatin occupancy and expands the repertoire of target genes

As mentioned earlier, HY5 over-expression is a major determinant of *det1-1* photomorphogenic phenotype (Pepper and Chory, 1997). Hence, to test whether HY5 excessive accumulation impacts on its activity at the chromatin level, we conducted ChIP-seq assays in WT and *det1-1* mutant seedlings grown under light conditions. To identify endogenous HY5 targets, we used an anti-HY5 antibody that gave no significant background (Fig. S2A) and was recently used for ChIP in Bellegarde *et al*., (2019). Additionally, *2×35S::GFP-HY5/hy5* (hereafter called *GFP-HY5*) functionally complemented seedlings were also used for an anti-GFP driven ChIP experiment to assess HY5 targets when the protein is overexpressed. GFP-HY5 level was higher than endogenous HY5 levels of WT and of *det1-1* seedlings (Fig. S2B). In our analysis, only genes with significant HY5 peaks in each of two independent biological replicates but not in the mock IP controls (*hy5-215* for the anti-HY5 ChIP and *2×35S::GFP* for anti-GFP) were considered as true HY5 binding genes for further analyses (Fig. S3A and B). This resulted in defining 422 HY5 target genes in WT seedlings (Fig. 5A), representing a smaller subset than in other studies reporting thousands of target genes (Kurihara *et al*., 2014; Hajdu *et al*., 2018). Among them, 372 genes (88%) were also identified in Hajdu *et al*., (2018) and/or by DAP-seq in O’Malley *et al*., (2016) (Fig. S3C) and 64 were common with the 297 HY5 so-called “directly-regulated genes” defined in Burko *et al*., (2020) (Fig. S3D).

**Figure 5.**
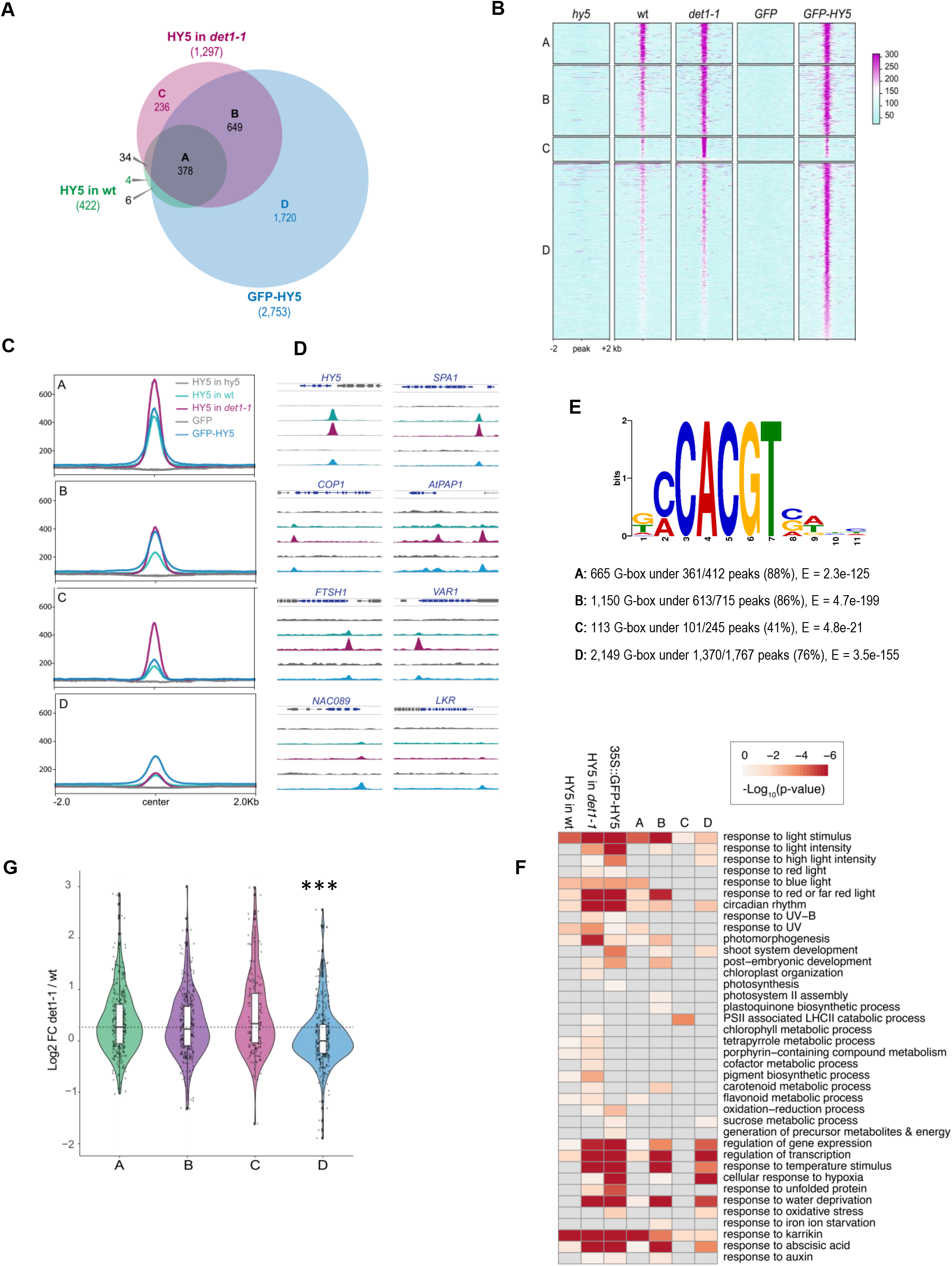
HY5 binds to extra-sites in *det1-1* background and in the GFP-HY5 overexpressor lines. (A) Comparisons of endogenous HY5 target genes in WT and *det1-1* and recombinant GFP-HY5 target genes in the overexpressing line. A represents the set of common target genes, B the extra HY5 target genes common to *det1-1* and GFP-HY5, while C and D the extra HY5 target genes found exclusively in *det1-1* or in the overexpression line, respectively. (B) Heatmaps showing the relative enrichment around the peaks found in the promoters of the A, B, C and D groups of genes. The HY5 IP in the *hy5* mutant background and the GFP IP in a line expressing an unfused GFP protein serve as mock controls. (C) Profiles showing the median enrichment around the peaks found in the promoters of the A, B, C and D groups of genes. (D) Snapshots of HY5 peaks on selected genes. (E) Enriched motifs were searched under the HY5 peaks found in the promoters of the A, B, C and D groups of genes. For each category the most highly enriched motif was a G-box/C-box. The number of occurrences of the motif and the E-value are stated below the motif logo. (F) GO enrichment analysis in the different group of genes. (G) The distribution of the Log2 fold change in expression in *det1-1* versus wild-type light-grown seedlings retrieved from Nassrallah *et al*. (2018) was plotted for the different gene categories. *** p-value<0.001.

Confirming the robustness of these endogenous HY5 genomic profiles, this HY5 target gene repertoire was also almost entirely conserved in the *det1-1* samples (only 10/422 target genes were not found in *det1-1*; Fig. 5A). In this mutant line, the HY5 targeted genes was extended to 1,297 (a 3-fold increase respect to WT). In line with a possible effect of HY5 over-accumulation in targeting extra sites, the number of HY5 associated genes further increased in GFP-HY5 plants to reach 2,753 genes (a 6.5-fold increase respect to WT).

We classified all gene sets in four groups, A representing the genes commonly bound in all three plant lines, B representing the genes significantly occupied in either *det1* or *GFP-HY5* (where HY5 over-accumulates), while C and D represent the genes specifically targeted in *det1-1* or in the *GFP-HY5* line, respectively (Fig. 5A). Remarkably, Class B represented a large set of 619 genes, indicating a robust tendency of HY5 binding on additional targets upon over-accumulation. Observation of these extra peaks in the WT sample unveils their initial pre-existence as low (but not statistically significant) ChIP signals, which get enriched to form robust peaks in the two plant lines with high HY5 global level (Fig. 5B, C and D). In line with these observations, HY5 occupancy also increases on class A (WT targets) in *det1-1* plants (Fig. 5C and D). These findings suggest that HY5 over-accumulation expands the HY5 cistrome, presumably as a consequence of TF rate-limiting availability that allows binding only to primary targets where its affinity is stronger or has better chromatin accessibility. Interestingly, HY5 global level on its own might not be the unique determinant of its DNA binding specificity, as HY5 enrichment over Class A genes is normal in *GFP-HY5* plants (Fig. 5C).

We therefore undertook a *de novo* sequence motif search, which identified that a CACGT motif is present under most of HY5 peaks in all gene classes but Class C (Fig. 5E). In this *det1-1* specific gene class, only 41% (versus 76-88% for A/B/D classes) of the peaks contain a G-box while, instead, several other DNA sequence motifs are over-represented in the extra-peaks identified in *det1-1* plants (Fig. S4).

To further test whether HY5 chromatin association is linked to gene expression, we first analysed the expression of the genes from classes A-D in *hy5-215* loss-of-function mutant plants using previously published RNA-seq (Myers *et al*., 2016) (Fig. S3E). Around 20% of the genes in each class were found to be misregulated, suggesting that only a minor proportion of HY5 target genes directly respond to HY5 occupancy. A majority of HY5 target genes may therefore be regulated by redundant transcription factors, as for example with the related HYH. As expected, gene classes A-B-C had a strong tendency for down-regulation in *hy5*, in agreement with HY5 being an activator of gene expression (Burko *et al*., 2020). Noteworthy, this was not the case for Class D, i.e., genes bound by HY5 only in the over-expressing *GFP-HY5* line (Fig. S3E).

To assess the effect of HY5 second-site binding on a genome-wide scale, we first determined the gene ontology (GO) of all different HY5 target gene sets. Extra-binding genes were found to be involved in the control of photosynthesis and pigment accumulation as well as to the response to stress responses as high light, temperature, UV-B, oxidative stress, hypoxia, water deprivation and nutrient deficit (Fig. 5F). Then, we analysed the expression of HY5-target genes in *det1-1* mutant seedlings using previously published RNA-seq (Nassrallah *et al*., 2018). Genes bound by HY5 in *det1-1* (Classes A, B and C) tend to be upregulated in *det1-1* mutant with respect to WT (Fig. 5G). Accordingly, HY5 target genes misregulated in *det1-1* are almost exclusively upregulated (Fig. S3F). On the contrary, class D genes (occupied by HY5 only in the overexpressed GFP-HY5 line) displayed tend to be equally expressed in WT and *det1-1* plants, confirming that this class of genes is specifically affected in the overexpressing line and not in *det1-1* (Fig. C). We therefore analysed the expression of a non-biased selection of class D genes in *GFP-HY5* seedlings, which showed that most of them are slightly down-regulated as compared to WT levels (Fig. S5). These observations support the possibility of an increasing repressive activity of HY5 when it over-accumulates. Misregulation of Class D genes and the identification of hundreds of secondary target genes indicate that Arabidopsis photomorphogenic seedlings require a tight modulation of HY5 levels, potentially avoiding out of context responses affecting multiple gene ontologies.

### HY5 and PIF3 targets overlap

DET1 has been found to associate with PIFs (Dong *et al*., 2014), that are thought to play antagonistic roles to HY5 in many aspects of plant development (Gangappa and Botto, 2016). This antagonism might rely on one hand on their differential regulation by light and on a potential competitive binding over a common gene repertoire bearing G-Box motifs as proposed earlier (Toledo-Ortiz *et al*., 2014; Gangappa and Kumar, 2017). To assess whether HY5 primary (WT) and secondary site (upon overexpression) binding may overlap the PIFs chromatin landscape, we performed PIF3 ChIP-seq experiments using a *pif3::eYFP:PIF3/pif3*-3 Arabidopsis line (Al-Sady *et al*., 2006) intended to mimic the endogenous PIF3 levels. Determination of PIF3 binding sites as done previously with HY5 identified 958 in dark-grown seedlings. In accordance with PIF3 being much more active under dark than under light conditions (Soy *et al*., 2012), most of the peaks called in light-grown plants were low intensity and correspond to genomic positions generating a high level of noise in the corresponding mock IPs and were therefore not considered as being robust for further analyses (Fig. 6A and B). PIF3-associated genes in darkness significantly overlap with a previous ChIP-seq experiment done on 2-day-old seedlings from an overexpression line and also with the repertoire of genes downregulated in the quadruple *pifQ* mutant line (Leivar *et al*., 2012; Zhang *et al*., 2013) impaired in the function of PIF1, PIF3, PIF4 and PIF5 partially redundant TFs (Fig. S6B and C). Confirming previous studies (Zhang *et al*., 2013), we detected a sequence motif matching the canonical motifs G-box (CACGTG) and PBE-box (CACATG) under half of PIF3 peaks (Fig. 6C). Two other over-represented motifs were also identified, with homology to TEOSINTE BRANCHED 1/CYCLOIDEA/PCF (TCPs) and, interestingly, FUS3 recognition motifs determined by DAP-seq (Fig. S6D; O’Malley *et al*., 2016). Altogether, these three motifs cover ~96% of the PIF3 peaks from our ChIP-seq, suggesting a high level of sequence specificity.

**Figure 6.**
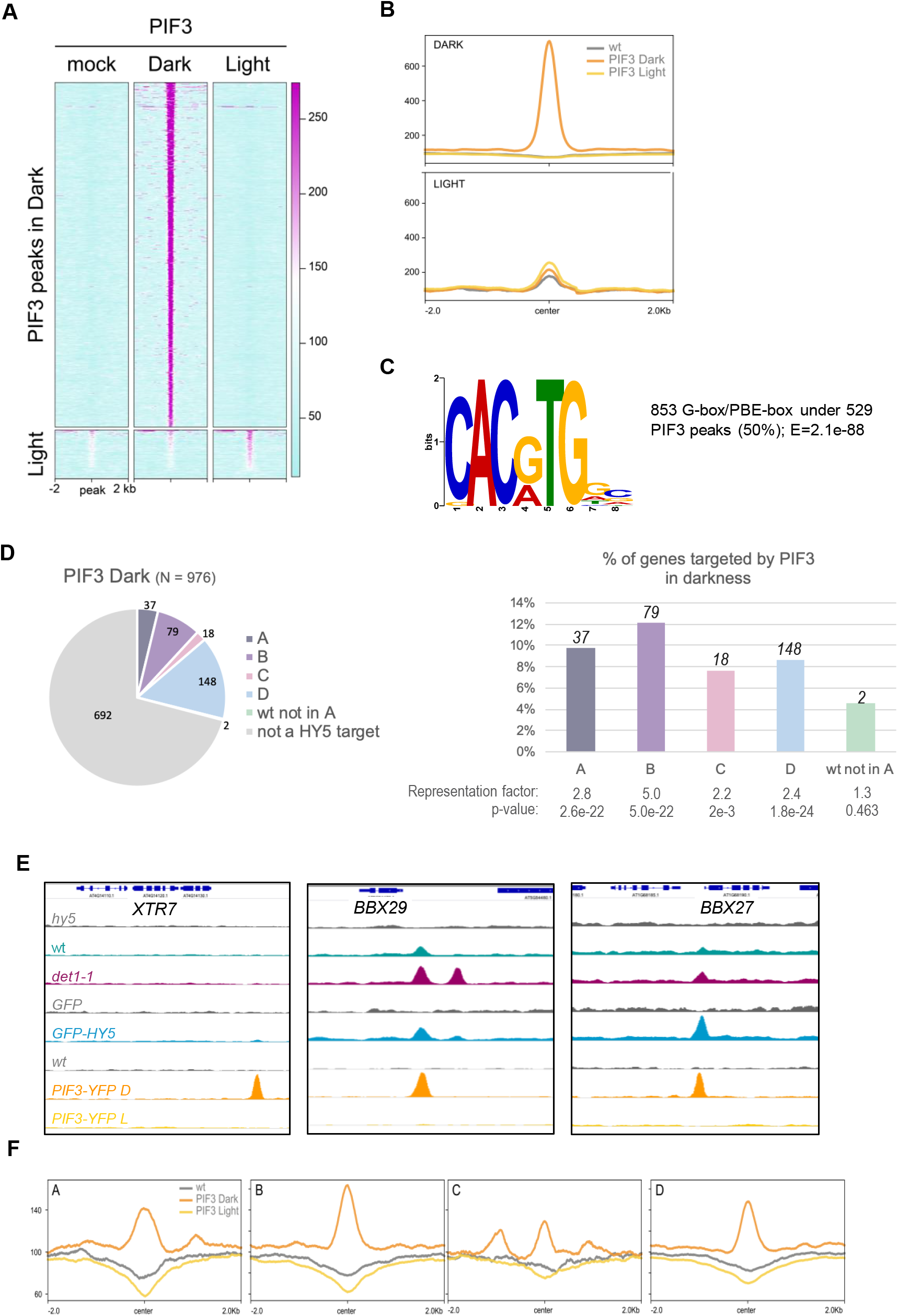
PIF3 targets in the dark display a partial overlapping with HY5 targets. (A) Heatmaps showing the relative PIF3 enrichment around the peaks found in dark and light conditions. (B) Profiles showing the median enrichment around the PIF3 peaks. (C) Enriched motifs were searched under the PIF3 peaks found in darkness. The most highly enriched motif matched the G-box/PBE-box. The number of occurrences of the motif and the E-value are stated next to the motif logo. (D) Left: proportion of genes belonging to the A, B, C and D classes among the PIF3 targets in darkness. Right: percentage of genes belonging to the A, B, C and D classes that are targeted by PIF3 in darkness. The significance of the enrichment is displayed below the graph. (E) Snapshots of HY5 and PIF3 peaks on 3 selected PIF3 target genes: *XTR7* is not targeted by HY5, *BBX29* belongs to class B and *BBX27* to class D. (F) Profiles showing the median PIF3 enrichment around the peaks found in the promoters of the A, B, C and D groups of genes.

Comparison of PIF3 with HY5 target genes restricts the shared repertoire to 39 loci that may consequently be simultaneously or alternatively occupied by both TFs in wild-type plants, representing ~4% of all PIF3 associated genes in darkness or ~9% of all HY5 associated genes in the light (Fig. 6D). Increased HY5 levels in *det1-1* and in the *GFP-HY5* lines resulted in an increased overlap with PIF3 target genes, with 79, 18 and 148 additional target genes found among classes B, C and D, respectively (Fig. 6D-F). This shows that HY5 secondary sites in the light tend to span loci occupied by PIF3 in darkness. Accordingly, analysis of PIF3 enrichment at HY5 peaks showed that HY5 and PIF3 peaks are centred around the same position, certainly corresponding to the G-box motif. A second PIF3 enrichment at positions neighbouring the main peak is detected for genes in class C, indicating that genes gaining HY5 in *det1-1* are frequently targeted by PIF3 at multiple positions in the promoter (Fig. 6F). Collectively, these analyses unveiled that an initial tendency for HY5 and PIF3 to target similar gene positions is exacerbated in *det1-1* and *GFP-HY5* plant lines, suggesting that moderation of HY5 accumulation in both light and dark conditions is required to avoid indirect effects on both HY5 and PIF3 regulatory networks.

### *HY5* over-expression causes *fusca*-like phenotype

Aiming at identifying potential out-of-context responses to HY5 over-expression, we analyzed the *GFP-HY5* complemented line used in our ChIP-seq analysis. Interestingly, attempts to obtain homogeneous complemented transgenic lines throughout generations were unsuccessful. Besides a variable number of non-germinated seeds, we obtained a segregating population of phenotypically “non-complemented” and “complemented” plants, with another group of small seedlings exhibiting typical “*fusca*-like” phenotypes of underdeveloped and purple plants with multiple growth defects including seedling lethality (Fig. 7A and B; Fig. S2B).

**Figure 7.**
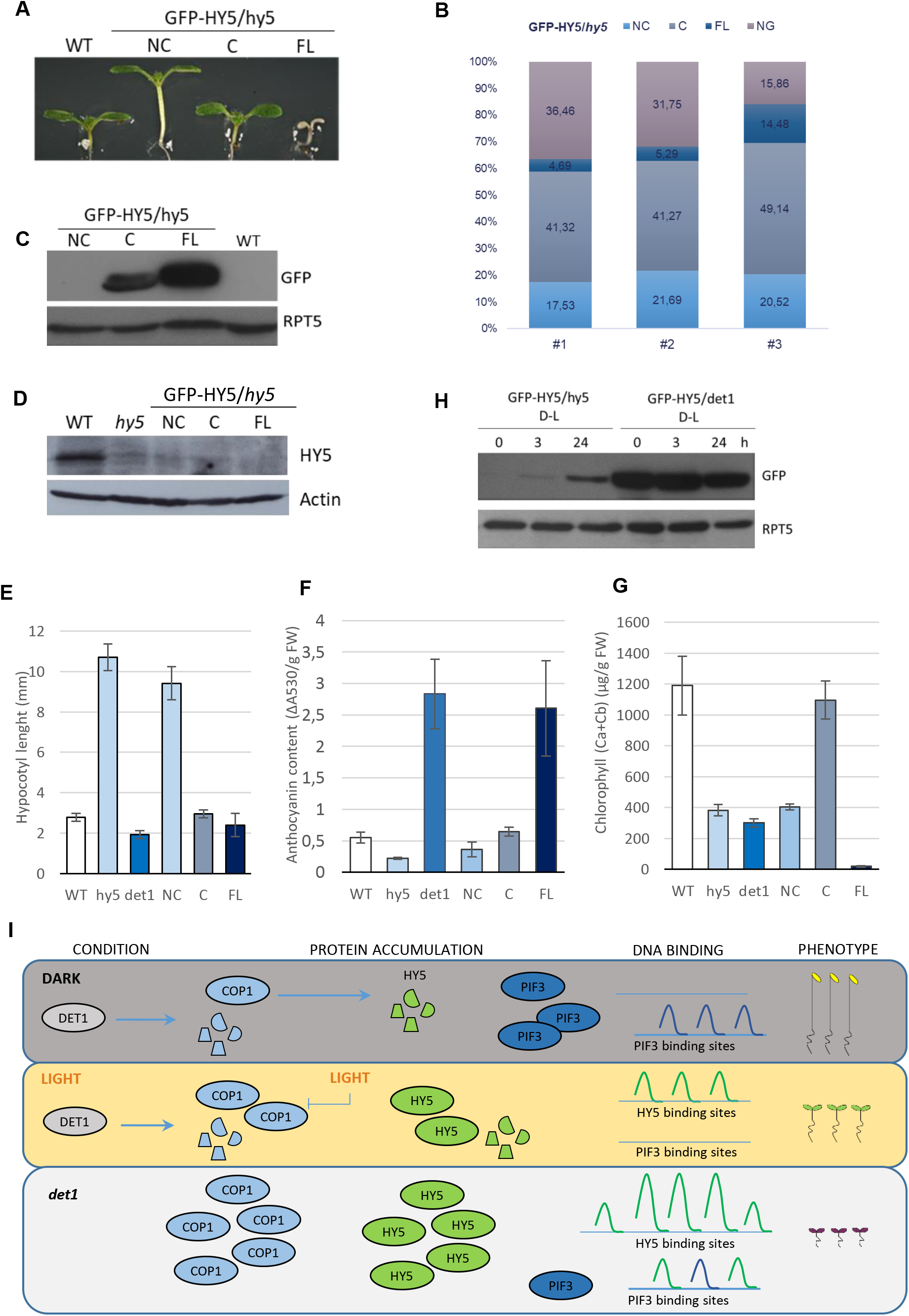
HY5 overexpression is sufficient to generate a *fusca*-like phenotype. (A) Seven day old seedlings representative of WT and the different phenotypes segregated from the GFP-HY5/*hy5* lines: non-complemented (NC), complemented (C, similar to WT) and *fusca*-like (FL) plants. (B) Frequency analysis of each phenotype on the segregating GFP-HY5/hy5 lines, and non-germinated seeds (NG). Lines #1, #2 and #3 are a result of independent transformation events. (C) GFP-HY5/*hy5* protein accumulation in the plants belonging to different types of GFP-HY5/*hy5* phenotypes. An anti-GFP was used to detect the GFP-HY5 fusion protein and an anti-RPT5 was used as a loading control. (D) Endogenous HY5 levels in seedlings displaying each of the GFP-HY5/*hy5* phenotypes, obtained by an immunoblot with anti-HY5 antibody showing that in all the plants endogenous HY5 is absent. Anti-Actin antibody was used for Actin loading control. (E) Hypocotyl measurements of the different phenotypical populations of GFP-HY5/*hy5* lines grown for 7 days under long day conditions. Bars represent average ± SD, n≥21. (F) Measurement of anthocyanin accumulation in 5 day old seedlings. Bars represent average ± SD of five measurements of pools containing a minimum of 15 seedlings. (G) Measurements of total chlorophyll (Ca+Cb) content of 7 day old seedlings. Bars represent average ± SD. (H) GFP-HY5 accumulation in GFP-HY5/*hy5* seedlings grown in the dark for 6 days and 3 or 24 hours after transfer to light. GFP-HY5 accumulation in *det1-1* background in the same conditions. An anti-GFP antibody was used and anti-RPT5 as loading control. (I) Working model for the role of DET1 on photomorphogenesis. By promoting COP1 protein destabilization, DET1 positively regulates COP1 activity towards the degradation of HY5. Keeping HY5 levels tightly regulated is essential to restrict its binding capacity (green peaks) over second-site targets. HY5 overaccumulation results in occupancy of new sites also targeted by PIF3 (blue peaks) in the dark. HY5 over-activity is sufficient to trigger *fusca*-like phenotypes and compromise plant viability.

Considering that appearance of this exaggerated photomorphogenic phenotype in several independent transgenic lines might represent the phenotypic translation of HY5 binding to an extended number of genes, we pursued the characterization of *fusca*-like plants. These seedlings accumulate greater levels of GFP-HY5 fusion protein than phenotypically complemented plants (Fig. 7C). Similarly to *fusca* mutants described in landmark studies from Misera *et al*., (1994) and Castle and Meinke genetic screenings (1994), *HY5*-based *fusca*-like plants display reduced growth, limited cotyledon expansion and high anthocyanin content especially in the cotyledons (Fig. 7E and F; S3C). They also present stronger defects in the aerial part of the plant as the root develops as in weak *fusca* mutants, perhaps correlating with a higher impact of HY5 activity in aerial tissues. Increased accumulation of anthocyanins occurs at early stages and relies on sugar as they do not accumulate when plants are grown in MS media without sucrose (Fig. S2D). This characteristic is also shared with original *fusca* mutant plants (Castle and Meinke, 1994) and further suggests an implication of HY5 in sugar induced anthocyanin accumulation.

Unlike original *fusca* mutant seedlings, HY5-based *fusca*-like plants turn pale across the days and present a strong defect on chlorophyll accumulation in the light, while behaving without significant phenotype when grown in darkness (Fig. 7G; Castle and Meinke, 1994; Misera *et al*., 1994). This correlates with GFP-HY5 protein level being undetectable by immunoblot analysis of wild-type etiolated seedlings and accumulating from 3 hours after transfer to light (Fig. 7H). This means that COP1 in the dark displays an enormous capacity to degrade HY5 protein present at much higher levels than those occurring endogenously and still, this capacity is completely impaired in a *det1-1* background (Fig. 7H).

Last, we examined expression of a selection of genes from clases A to D in light-grown *fusca*-like seedlings to test whether such phenotype can be linked to HY5 association to chromatin secondary targets. This unveiled stronger misregulation in this category of plants than in functionally complemented *GFP-HY5* plants. While in some cases the *fusca*-like phenotype can be associated to higher gene upregulation (as in the case of *CHS, F3H, CRF6, CUC1* or *WOX1*), in the majority of genes tested they were associated to a repressive effect suggesting that HY5 extra occupancy contributes to triggering *fusca*-like phenotypes. Interestingly, this feature is true for the majority of PIF3 direct targets that are also targeted by HY5 (e.g., *BBX28, TCP2, RGA2/GAI, GASA6, BBX27, XTH7*).

Altogether, these results provide evidence that exaggerated HY5 accumulation by itself is sufficient to generate *fusca*-like phenotypes and render plants unviable under light conditions. At several loci, HY5-derived gene repression occurs through ectopic enrichment over secondary sites, potentially at the expenses of PIF3 occupancy over a set of commonly targeted genes.

## Discussion

DET1 and COP1 were identified more than 30 years ago as repressors of photomorphogenesis and of multiple light responses. Impairment of any of these proteins results in the induction of deetiolation in darkness and in hyper-photomorphogenic or so-called *fusca* phenotypes in the light. Because they form CRL4 based E3 ligases involved in HY5 degradation, COP1 and DET1 were expected to work in a close relationship whose nature was never elucidated. We identified that COP1 and DET1 can associate–together, DET1 controlling COP1 stability by promoting its proteasomal degradation which seems to be essential for COP1 activation.

DET1-mediated regulation of COP1 is of prime importance given its necessity for the regulation of HY5 abundance, a feature that seemingly look paradoxical as *det1* mutant plants over-accumulate both COP1 and HY5 proteins. In view of these observations, we propose that DET1-mediated COP1 destabilization is necessary to maintain COP1 turnover and activity towards its downstream targets. We also unveiled that, by down-regulating HY5 levels, DET1 restrains HY5 binding to primary targets. Over-expression of HY5 can largely extend its cistrome, not only by inducing HY5 enrichment over second-sites (normally poorly bound) target genes but also by occupying PIF3 target genes. As detailed below, HY5 second-site occupancy correspond to multiple categories of light-regulated genes and was further found to be linked in *cis* to gene misregulation, a property that presumably underlies the frequent occurrence of *fusca*-like phenotype upon artificial *HY5* over-expression (Fig. 7I).

### DET1 conforms CRL4^C3D^ complexes and associates with CSN and with COP1

Together with DDB1, COP10 and DDA1, DET1 forms a stable C3D complex that associates with the CUL4 scaffold as well as with the CSN (Schroeder *et al*., 2002; Wertz *et al*., 2004; Yanagawa *et al*., 2004; Chen *et al*., 2006; Olma *et al*., 2009; Lau and Deng, 2012; Irigoyen *et al*., 2014). Our TAP assays showed that most of DET1 engages in the formation of the C3D complex and only part of this complex associates with CUL4. A smaller fraction of DET1, perhaps in the form of CRL4^C3D^ complexes, associated with a fully assembled CSN. CSN-associated CRL4^C3D^ might represent the substrate-free fraction, since substrate binding to CRL4 is expected to displace the CSN (Cavadini *et al*., 2016). Moreover a set of WD40 domain containing proteins, known as typical CRL4-associated target receptors, also co-purified with DET1 (Table S3; Fonseca and Rubio, 2019). Among them, we found COP1 and SPA1 proteins and reciprocally, DET1 peptides were found in TAP assays when COP1 was used as a bait. By means of co-immunoprecipitation assays, previous reports discarded a DET1-COP1 association in Arabidopsis even though COP1 and the C3D subunit COP10 were found to interact, presumably independently of the C3D complex (Chen *et al*., 2010). This supported the idea that DET1 and COP1 exist in distinct complexes, each of them conforming CUL4 based E3 ligases, CRL^C3D^ and CRL4^COP1SPA^ (Schroeder *et al*., 2002; Yanagawa *et al*., 2004; Chen *et al*., 2010) and that they may work together, repressing photomorphogenesis in an unsolved way (Lau and Deng, 2012). For the first time, we demonstrate here that, similar to human cells (Wertz *et al*., 2004), DET1 and COP1-HY5 module can associate in Arabidopsis.

As previously reported, we also detected that COP1 further associates with all four SPA proteins (specially with SPA4) and with CRY2 (and CRY1 with less affinity) (Wang *et al*., 2001; Yang *et al*., 2001; Saijo *et al*., 2003; Laubinger *et al*., 2004; Liu *et al*., 2011). It was however surprising that we could not recover CUL4 or CSN peptides from COP1 TAPs, as COP1 has been described to associate with CUL4 to stably form CRL4^COP1SPA^ complexes (Chen *et al*., 2010). This might be due to transient association of COP1 with CUL4 or to limited resolution capacity of our MS analysis; which might be insufficient to capture the whole catalogue of complexes conformed by COP1.

### DET1 promotes COP1 degradation and activity

Following the identification of a DET1-COP1 association, our study sheds light on the nature of this relationship, DET1 being necessary for COP1 protein destabilization. COP1 accumulates at higher levels in the light than under dark and is a short-lived protein with a fast turnover rate in both conditions (estimated half-life of 4,5 hours, Fig. S1B). DET1-mediated COP1 degradation is light independent and depends at least in part on the proteasome. Our mutants’ analyses indicate that COP1 degradation mechanism seems to rely on a canonical CSN-mediated CUL4 recycling that is mediated by a full C3D complex.

Our biochemical findings seemingly enter in conflict with genetics, because *det1-1* seedlings, as well as *cop1-4*, are deetiolated in the dark, meaning that DET1 and COP1 generally repress light signalling. Accordingly, for this reason, we wished to confirm that, as previously reported, HY5 levels are higher in both mutant backgrounds (Fig. 4A; Osterlund *et al*., 2000). In 1994, Ang and Deng analysed the epistatic relationships between *cop1* and *det1* hypomorphic mutations. They found *cop1-6* is epistatic to *det1-1*, with respect to light control of seed germination and dark-induced gene expression, suggesting that DET1 and COP1 may act in the same pathway, with COP1 being downstream, which fully supports our findings.

In our study, COP1 protein levels do not simply correlate with its activity when using its capacity to degrade HY5 as readout. Indeed, low levels of COP1 in darkness are sufficient to degrade both physiological and over-accumulated HY5 protein levels, as those displayed by wild-type and *GFP-HY5* overexpressing lines, respectively (Fig. 7H). Counter-intuitively, higher COP1 protein levels following exposure to light or in the *det1-1* mutant are associated to diminished COP1 function. This effect might result from COP1 activity being regulated by its association with SPA proteins or photoreceptors and also through nucleo-cytoplasmic partitioning (Wang *et al*., 2001; Yang *et al*., 2001; Laubinger and Hoecker, 2003; Seo *et al*., 2003; Laubinger *et al*., 2004; Fankhauser and Ulm, 2011; Lian *et al*., 2011; Liu *et al*., 2011; Rizzini *et al*., 2011; Zuo *et al*., 2011; Ponnu *et al*., 2019). We found a large amount of COP1 in the nuclei of *det1-1* mutants, precluding any inactivation of the COP1 protein pool through nuclear exclusion. Alternative molecular mechanisms should be envisaged, as for example DET1 promotion of COP1 association with SPA proteins or any other member of CRL4^COP1-SPA^ complexes. As well, DET1 could promote CSN-mediated cycles of CUL4 neddylation/deneddyation required for CRL4^COP1-SPA^ activity. Finally, following on our results showing DET1-HY5 coprecipitation, DET1 could facilitate substrate recognition and ubiquitination by COP1. All these processes might be necessary for efficient ubiquitination and proteasomal degradation of the COP1 substrates but also of COP1 itself, as a feedback mechanism to limit the extent of its activity.

### Higher HY5 accumulation increases binding to extra targets including PIF3 target genes

HY5 is a pivotal TF in light signalling with a strong effect on plant morphogenesis, whose levels are tightly regulated (Osterlund *et al*., 2000). Therefore, the identification of its direct genomic targets to identify genes directly regulated by HY5 fundamentally needs to be considered in the context of dynamic changes in HY5 global level in the nucleus. In other words, as proposed for PIF transcription factors (Pfeiffer *et al*., 2014), HY5 chromatin association and its sets of targeted genes need to be envisaged as a potential continuum that varies with HY5 protein availability, chromatin accessibility and the abundance of other TFs potentially binding competitively to the same loci. Adding to technical variability, this concept might be central in the large variations of HY5 target genes from previous studies that reported lists reaching ~12,000 HY5 targets genes (Lee *et al*., 2007; Zhang *et al*., 2011; Kurihara *et al*., 2014; Hajdu *et al*., 2018). Probing endogenous HY5 protein, our ChIP analysis led to the identification of a moderate number of 422 targets that largely overlap the repertoire of 297 high-confidence HY5-activated genes reported by Burko and co-workers (2020) using an elegant strategy combining transcriptional and ChIP analyses of constitutive activator and repressor HY5 fusion proteins. Still in line with Burko *et al*., (2020), we found that in WT in light conditions HY5 behaves mainly as a transcriptional activator. Among HY5 targets in light-grown seedlings, we found previously described light-regulated HY5-bound genes such as the *HY5* gene itself and many other genes with the capacity to trigger downstream transcriptional cascades influencing a range of light-regulated processes: light stress (*ELIP1*), pigment biosynthesis (*CHS, F3H, FLS1*), signalling proteins (*SPA1, SPA3* and *SPA4*) as well as a high number of transcription factors (Table S4) (Oyama *et al*., 1997; Gangappa and Botto, 2016). HY5 peak summits were positioned on a typical or a related G-box sequence motif in the vast majority of these genes, but traces of HY5 binding could also be found over many other loci, potentially secondary or cell-specific. When identifying that many of these second-site binding loci are increasingly occupied by HY5 in *det1-1* and in the *GFP-HY5* over-expression line hints at the necessity for HY5 level fine-tuning to restrict the activity of this TF over a specific set of loci. This also hints at potential variations of the HY5 target gene repertoire during dark-to-light or light-to-dark transitions when HY5 abundance (Osterlund *et al*., 2000) and chromatin properties (Bourbousse *et al*., 2020) are subjected to strong variations.

Considering the preponderant role of HY5 over-accumulation in *det1-1* photomorphogenic phenotypes (Pepper and Chory, 1997), HY5 enrichment over second-site target genes likely contributes to gene misregulation induced by *DET1* loss of function. For example, among the genes significantly upregulated and targeted by HY5 specifically in *det1-1* plants (class C) (Fig. S3F), we identified four subunits of the chloroplast FtsH protease complex (FTSH1, 2, 5 and 8 found in the GO category “PSII associated LHCII catabolic process”; Fig. 5F) involved in the quality control of the photosynthetic electron transfer chain during photo-oxidative stress (Kato and Sakamoto, 2018). Misregulation of such stress response pathways must have a high cost impact on plant growth and performance as observed in *det1* plants.

Gene misregulation of HY5 secondary targets in *det1-1* and GFP-HY5 plant lines might result from a combination of multiple mechanisms, ranging from HY5 capacity to activate transcriptional, ectopic recruitment of chromatin machineries such as the GCN5 acetyltransferase (Benhamed *et al*., 2006) and, among other effects, competitive binding with other TFs. Comparison of our HY5 and PIF3 ChIP experiments indicate that HY5 enrichment over additional targets leads to a large increase in the overlap with PIF target genes, as for example 245 genes occupied by PIF3 in the dark are newly bound by HY5 when in *det1-1* and/or *GFP-HY5* overexpressor (Fig. 6D). Reduced abundance of PIFs in *det1-1* mutant plants (Dong *et al*., 2014) might facilitate HY5 binding to common targets with PIF3, but this is presumably not the case in the *GFP-HY5* overexpressing line. Expression analyses indicate genes downregulated in *det1-1* and GFP-HY5 lines are PIF3 targets, thereby suggesting that HY5 enrichment is translated into a higher repressive activity (Fig. S5).

Taken together, these interplays indicate that TFs availability and balanced levels are key to account for differential binding of HY5 and PIF3 proteins. It has been previously proposed that HY5 could compete with PIFs for binding sites on DNA for specific gene targets. For instance, HY5 and PIF4 proteins bind with different intensities to common targets genes at different day-times and temperatures (Toledo-Ortiz *et al*., 2014; Gangappa and Kumar, 2017). This idea is fully supported by our data in a genome-wide context. In future studies, it will be interesting to test HY5 binding specificity in *pif* higher-order mutant plants and, *vice-versa*, to assess the influence of HY5 on PIF chromatin landscape when their respective levels are balanced during dark-light transitions.

### Uncontrolled HY5 accumulation triggers *fusca*-like phenotypes

At high levels, HY5 chromatin association exceeds its primary target genes to an increased number of target sites, with consequent transcriptional changes. In line with the concept of fine-tuning TF abundance, appearance of *fusca*-like phenotypes in *GFP-HY5* plants show the pleiotropic effects derived from the uncontrolled accumulation of a single transcription factor, especially when, like HY5, it regulates many signalling pathways and other TFs. The *fusca* mutant plants isolated in the 90’s (Castle and Meinke, 1994; Misera *et al*., 1994) accumulate high levels of anthocyanin in the seeds and seedlings, display light independent (constitutive) seed development and compromised viability. These *fusca* mutants were shown to be impaired in COP1, DET1 or CSN activity, which all contribute to moderate HY5 accumulation, a process probably enhanced by a *cis*-acting positive feedback loop linked to HY5 autoactivation (Chory *et al*., 1989; Deng *et al*., 1991; Mayer *et al*., 1996). Across the years however, studies based on the transcriptional analysis of the *fusca* mutants suggested that these phenotypes could not be supported uniquely by altered light responses and should be due to a general defect in developmental programming, because several other signal transduction pathways were affected. These pathways described by Mayer *et al*., (1996), widely overlap with those present in the GO analysis of HY5 target gene classes B, C and D (Fig. 5F), showing that they are spanned by HY5 action. Gene expression analysis of key transcription factors involved in developmental processes such as cytokinin signalling (CRF6), meristem maintenance and initial organ development (WOX1, CUC1) and circadian clock (TOC1) showed these genes are upregulated in *fusca*-like plants (Fig. S5; Takada *et al*., 2001; Dolzblasz *et al*., 2016; Kim, 2016; Fung-Uceda *et al*., 2018). Reciprocally, a number of TFs (e.g. *CBF3, BBX28, TCP2, HFR1, BBX27*) and photosynthesis related genes (*LHCA1, LHCB7, FTSH1, FTSH5, PAP2, PSAE*) are downregulated in *fusca*-like seedlings. For the latter ones, HY5 apparently behaves as a transcriptional repressor by occupying extra target sites shared with PIF3 in darkness (Fig. 6D). Through this mechanism, HY5 may control numerous processes necessary for plant viability, including meristem activity, cell cycle, pigment accumulation and photoautotrophy.

Thus, in line with the finding that it lacks its own activation or repression domains (Ang *et al*., 1998), HY5 transcriptional regulatory activity over a limited gene repertoire might be regulated by titration of its availability. COP1 and DET1 activities are part of this regulatory mechanism to keep HY5 transcriptional activation sharp and responsive to light perception in a dynamic system.

## Material and Methods

### Plant Materials

All plant lines are in the Columbia-0 ecotype background. The *det1-1* (Chory *et al*., 1989), *hy5-215* (Osterlund *et al*., 2000), *det1-1hy5-215* was kindly provided by Prof. Roman Ulm (Geneva, Switzerland); *cop1-4* (McNellis *et al*., 1994), *pif3::eYFP:PIF3/pif3-3* (Al-Sady *et al*., 2006); kindly provided by Drs Lot Gommers and Elena Monte, CRAG Barcelona, Spain). The *2×35S::GFP-HY5/hy5-215* was generated by *Agrobacterium tumefaciens* (GV3101) and floral dip (Clough and Bent, 1998) transformation of *hy5-215* mutants with a GFP-HY5 expressing plasmid based on the pVR TAP Nt plasmid where the TAP tag cassette was substituted by the GFP reporter gene (Rubio and Deng, 2008).

### Plant growing conditions

Arabidopsis seedlings were sterilized with a solution of 75% sodium hypochlorite and 0.1% Tween-20 and stratified at 4°C during 3 days in darkness. Seedlings were grown in Murashige and Skoog (MS) medium with 1% sucrose and 0.7% agar at 22°C for 7 days (unless otherwise specified) under fluorescent white light (100 μmol m^-2^ s^-1^) in a 16-h light/8-h dark period (LD).

### Hypocotyl measurements

For hypocotyl measurements 7 days-old plants were disposed on agar plates, photographed and hypocotyls measured using ImageJ software (http://www.imagej.net). Three biological replicates, each consisting of measurements for at least 30 seedlings grown at different times, were analyzed with similar results.

### Protein extraction and immunoblotting

In the indicated experiments, 6 to 7 day old light or dark grown seedlings were pre-treated with 50 μM cycloheximide (CHX, Sigma Aldrich) or with 50 μM proteasome inhibitor Bortezomib (Selleckchem). For COP1 detection, extraction of plant soluble protein extracts was performed in 4 M Urea, 50 mM Tris-HCl, pH 7.4, 150 mM NaCl, 10 mM MgCl_2_, 1 mM phenylmethylsulfonyl fluoride, 0.1% Nonidet P-40 and cOmplete EDTA-free protease inhibitor cocktail (Roche) supplemented with followed by centrifugation twice 10 min at 16,000 g at 4°C. Protein concentration in the final supernatants was determined using the Bio-Rad Protein Assay kit. For HY5 detection, an equal number of plants were collected for each condition, directly denatured in Laemmli buffer and separated by a 15% blotted with antibodies described below. Chromatin-enriched protein fractions were obtained as previously described (Nassrallah *et al*., 2018).

### Pull-down assays

MBP recombinant protein fusions were expressed in the *Escherichia coli* BL21 (DE3) strain carrying the corresponding coding sequence cloned into the pKM596 plasmid, a gift from David Waugh (Addgene plasmid # 8837). Recombinant proteins were purified and pull-down assays were performed according to Fonseca and Solano, (2013). MBP-tagged fusions were purified using amylose agarose beads. Equal amounts of seedling protein extracts were combined with 10 mg MBP-tagged fusion or MBP protein alone, bound to amylose resin for 1 hr at 4°C with rotation, washed three times with 1 ml of extraction buffer, eluted and denatured in sample buffer before immunoblot analysis.

### TAP Assays

Cloning of a GSRhino-TAP–tagged DET1 and COP1 fusion under the control of the constitutive cauliflower mosaic virus 35S promoter, transformation of PSB-D Arabidopsis cell suspension cultures and TAP purifications were performed as described previously (Van Leene *et al*., 2015; García-León *et al*., 2018). For the protocols of proteolysis and peptide isolation, acquisition of mass spectra by a 4800 MALDI TOF/TOF Proteomics Analyzer (AB SCIEX), and mass spectrometry–based protein homology identification based on the TAIR10 genomic database, we referred to Van Leene *et al*., (2010) and García-León *et al*., (2018). Experimental background proteins were subtracted based on 40 TAP experiments on wild-type cultures and cultures expressing TAP-tagged mock proteins GUS, RFP, and GFP (Van Leene *et al*., 2010).

### ChIP analyses

HY5 ChIPs were performed on 5 day-old seedlings grown under LD conditions. PIF3 ChIPs were performed on 5-day-old seedlings grown in darkness or LD conditions. ChIP experiments were performed as in Fiorucci, *et al*. (2019), using anti-HY5 (Agrisera #AS121867) or anti-GFP (Life Technologies #11122).

#### Library preparation, sequencing, and analysis

Libraries were prepared using the NEBNext Ultra II DNA Library Prep Kit (New England Biolabs E7645). Sequencing was performed on an Illumina HiSEq 4000 in 150-bp paired-end mode. Reads were trimmed using trim_galore (https://github.com/FelixKrueger/TrimGalore) with options “--phred33 --paired -q 20 --stringency 1 --length 35” and then mapped to the TAIR10 genome using bowtie2 (Langmead and Salzberg, 2012) with options “--very-sensitive -I 150 -X 2000 -p 20 --no-mixed”. Duplicated reads were marked using picard-tools (https://github.com/broadinstitute/picard). Reads were then filtered using samtools (https://github.com/samtools/samtools.git) with options “view -hb -F 1804 –L selected_TAIR10_genome_Chr.bed”, which corresponds to the TAIR10 genome after filtering out genomic regions with aberrant coverage or low sequence complexity (Quadrana *et al*., 2016). Browser tracks were generated using deeptools (Ramírez *et al*., 2016) function bamCoverage with options “--binSize 20 --normalizeUsingRPKM --extendReads --centerReads --ignoreForNormalization ChrC ChrM”. Peaks were called using MACS2 (McNellis *et al*., 1994) with options “callpeak -f BAMPE --bdg -q 0.01 -g 120e6 --bw 300” and using the input bam files as control. The peaks present in the mock IPs were removed for further analysis using bedtools (Quinlan and Hall, 2010) with options “subtract -A -f 0.2” and only the remaining peaks with a score above 60 were kept. A second filtering step consisted in keeping only the peaks present in both biological replicates using bedtools with options “intersect -f 0.2 -r”. Peaks were then annotated to the closest TSS using HOMER annotatePeaks.pl (Heinz *et al*., 2010) providing Araport11 gtf annotation file (Cheng *et al*., 2017). Heatmap and metaprofiles were generated using deeptools computeMatrix, plotHeatmap and plotProfile functions.

#### Motif and GO enrichment search

Motif search was performed using meme from the MEME suite (Bailey *et al*., 2009) with options “-dna -mod anr -revcomp -maxsize 25000000 -nmotifs 10 -minw 6 -maxw 12 -maxsites 10000 -brief 3000 -p 5”. All found motifs were then compared with the DAP-seq database of motifs (O’Malley *et al*., 2016) using Tomtom (Gupta *et al*., 2007) with default options. The Gene Ontology enrichment analyses were performed using GO-TermFinder (Boyle *et al*., 2004) via the Princeton GO-TermFinder interface (https://go.princeton.edu/cgi-bin/GOTermFinder), and then simplified using REVIGO (Supek *et al*., 2011; Langmead and Salzberg, 2012) and visualized as an unclustered heatmap using pheatmap (https://cran.r-project.org/package=pheatmap).

### Pigment quantification

For anthocyanin quantification the aerial parts of 15 to 20 5-day-old seedlings, collected from different plates, were pooled for each replicate. Anthocyanin quantification was performed as described in Hillis and Swain, 1959. Six to 10 7-day-old seedlings were pooled for chlorophyll measurements. Acetone 80% (V/V) was used for extraction and A645 and A663 was measured in a spectrophotometer Data analysis was done according to Arnon, 1949. Three independent replicates (seedling pools) were measured for each sample. Values represent mean ± SD.

### RNA analyses

For RT-qPCR assays, 2 ug total RNA extracted from 7 day old seedlings with the Favorprep Plant Total RNA Purification Mini kit (Favorgen) was used for cDNAs synthesis with using the High-Capacity cDNA Reverse Transcription kit (Applied Biosystems) with DNAse I treatment (Roche). Quantitative PCR was carried out using 5x PyroTaq qPCR mix Plus EvaGreen (CMB Cultek Molecular Bioline) in a QuantStudio5 machine (Applied Biosystems). Transcripts were amplified and results were normalized to *PP2A* transcript levels. Primers used for QPCR are represented in Table S5.

### Antibodies

Antibodies used for immunoblot experiments: anti-GFP-HRP (Milteny Biotec #130-091-833); anti-H2B (Millipore #07–371); anti-MBP (Abcam #9084); anti-Actin (Sigma #A04080); anti-RPT5 (Enzo Life Sciences# BML-PW8245); anti-HY5 (Agrisera #AS121867 or Abiocode #R1245-1b); anti-COP1 (kindly provided by Xing Wang Deng); anti-mouse (ThermoFisher Scientific #A11001) or anti-rabbit (ThermoFisher Scientific #A11008) secondary antibodies.

### GEO accession

ChIP-seq data generated in this work are accessible through GEO Series accession number GSE155147.

## Author Contributions

C.B., F.B., V.R and S.F., designed the research and conceived the study. E.C., C.B., M.G-L., L.W., C. G-B., V.R. and S.F. performed experiments. C.B. analysed ChIP-seq data. C.B., F.B., V.R. and S.F. discussed the results; C.B. and S.F. wrote the initial version of the paper and V.R. and F.B. edited the manuscript.

## Acknowledgements

We are grateful to Roberto Solano and Salomé Prat for the critical reading and suggestions on the manuscript.

## Supplemental Information

Supplemental files contain:

**Figure S1. COP1 expression levels in different mutants.**

**Figure S2. GFP-HY5/*hy5* line analysis.**

**Figure S3. Analysis of HY5 targets in comparison with previous published binding and expression data.**

**Figure S4. *De novo* motif search under HY5 peaks annotated A to D gene classes.**

**Figure S5. Gene expression analysis of HY5 bound genes.**

**Figure S6. Analysis of PIF3 targets expression and binding sites and overlapping with HY5 binding classes.**

**Table S1. Interactomics of DET1 and COP1 proteins.**

**Table S2. List of DET1 and COP1 associated proteins found in the different replicates of TAP assays.**

**Table S3. List of DWD proteins that associate with DET1.**

**Table S4. Transcription factors that are direct targets of HY5 in wild-type.**

**Table S5. Oligonucleotides used in this study.**

**Supplemental Data 1 - HY5 targeted genes.**

**Supplemental Data 2 - PIF3 targeted genes in the dark.**

