## Supplementary information for "DET1-mediated COP1 regulation avoids HY5 activity over second-site targets to tune plant photomorphogenesis"

Supplemental information

**A**

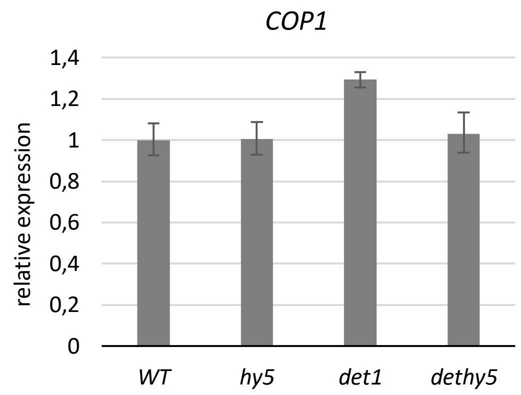

**B**

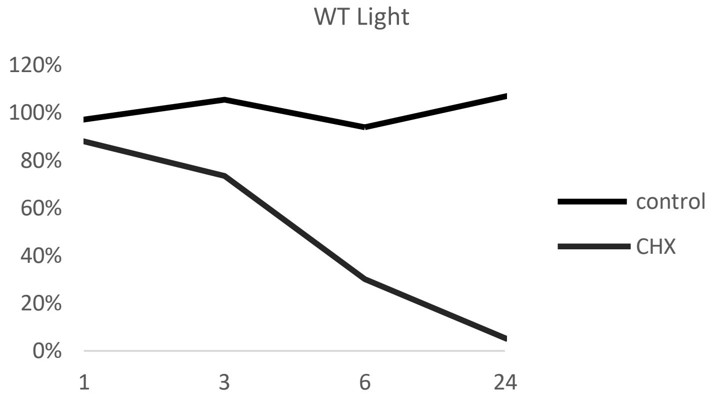

**Figure S1. COP1 expression levels in different mutants.**
(A) Analysis of *COP1* transcript levels in *hy5-215*, *det1-1*, and *det1-1hy5-215* mutants. The level of *COP1* in WT light-grown seedlings was set as 1. *PP2A* levels were used as endogenous controls. Bars represent average ± SD of three technical replicates.
(B) Arabidopsis COP1 degradation rate in white light conditions. Estimated half life of 4,5 hours. The graphic is based on protein quantification of Fig. 2E with ImageJ software. Band intensity was quantified respect to RPT5 in each sample and time 1 hour from control was taken as 100%.

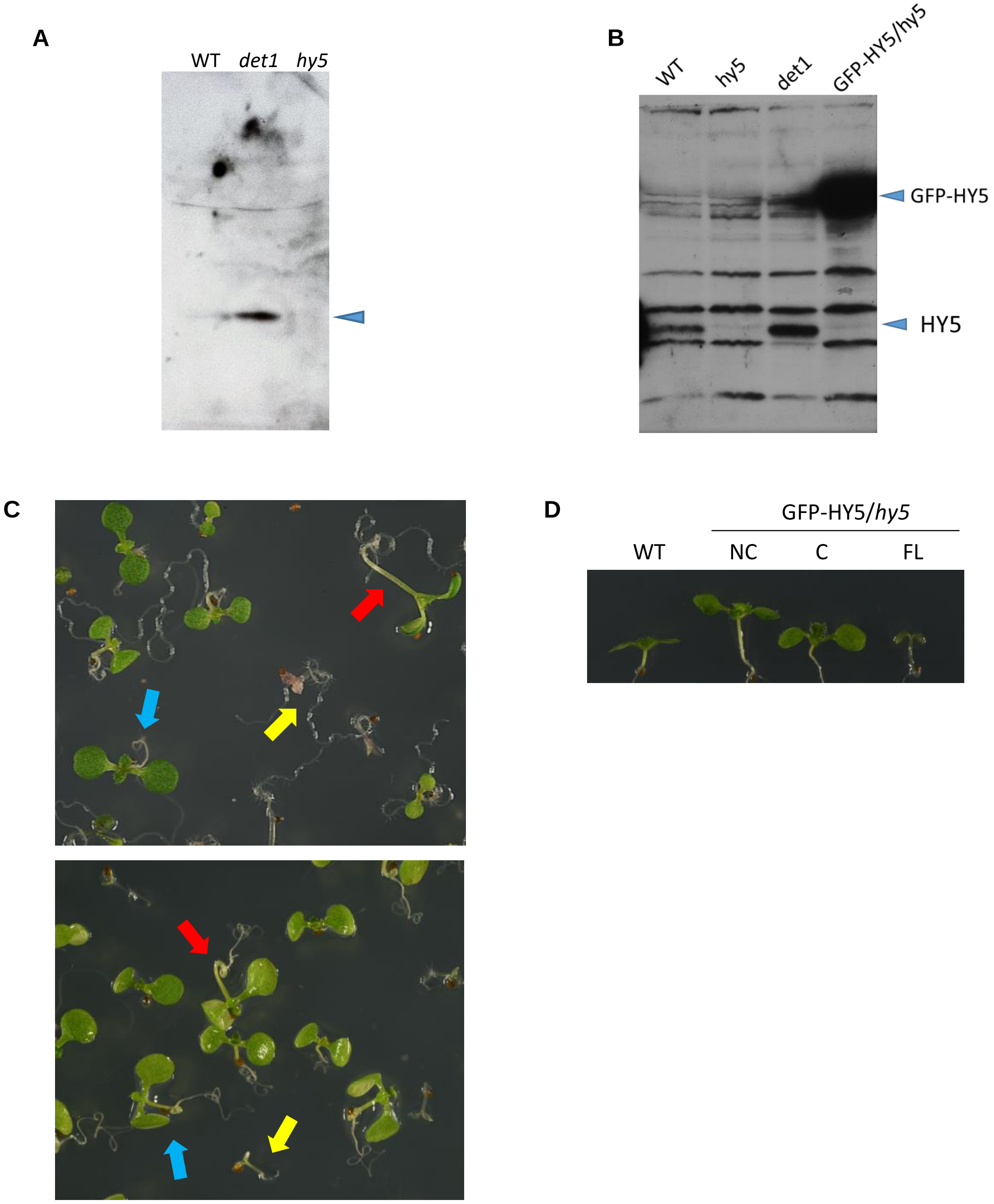

**Figure S2. GFP-HY5/*hy5* overexpression line analysis.**
(A) Specificity of the anti-HY5 antibody (Agrisera) used for ChIP experiments. Total protein extracts loaded in a 15% SDS page. The arrow indicates HY5 protein.
(B) HY5 accumulation levels in GFP-HY5 complemented plants and the endogenous HY5 protein levels detected with an anti-HY5 antibody (Abiocode).
(C) GFP-HY5/*hy5* seedlings grown in, Up: MS with 1% sucrose, Down: MS without sucrose, under long-day conditions. Complemented (blue arrow), non-complemented (red arrow) and *fusca*-like (yellow arrow) plants are visible.
(D) Seven day old seedlings grown in MS without sucrose under LD conditions. Representative seedlings of WT and the different phenotypes segregated from the GFP-HY5/*hy5* lines: non-complemented (NC), complemented (C, similar to WT) and *fusca*-like (FL) plants.

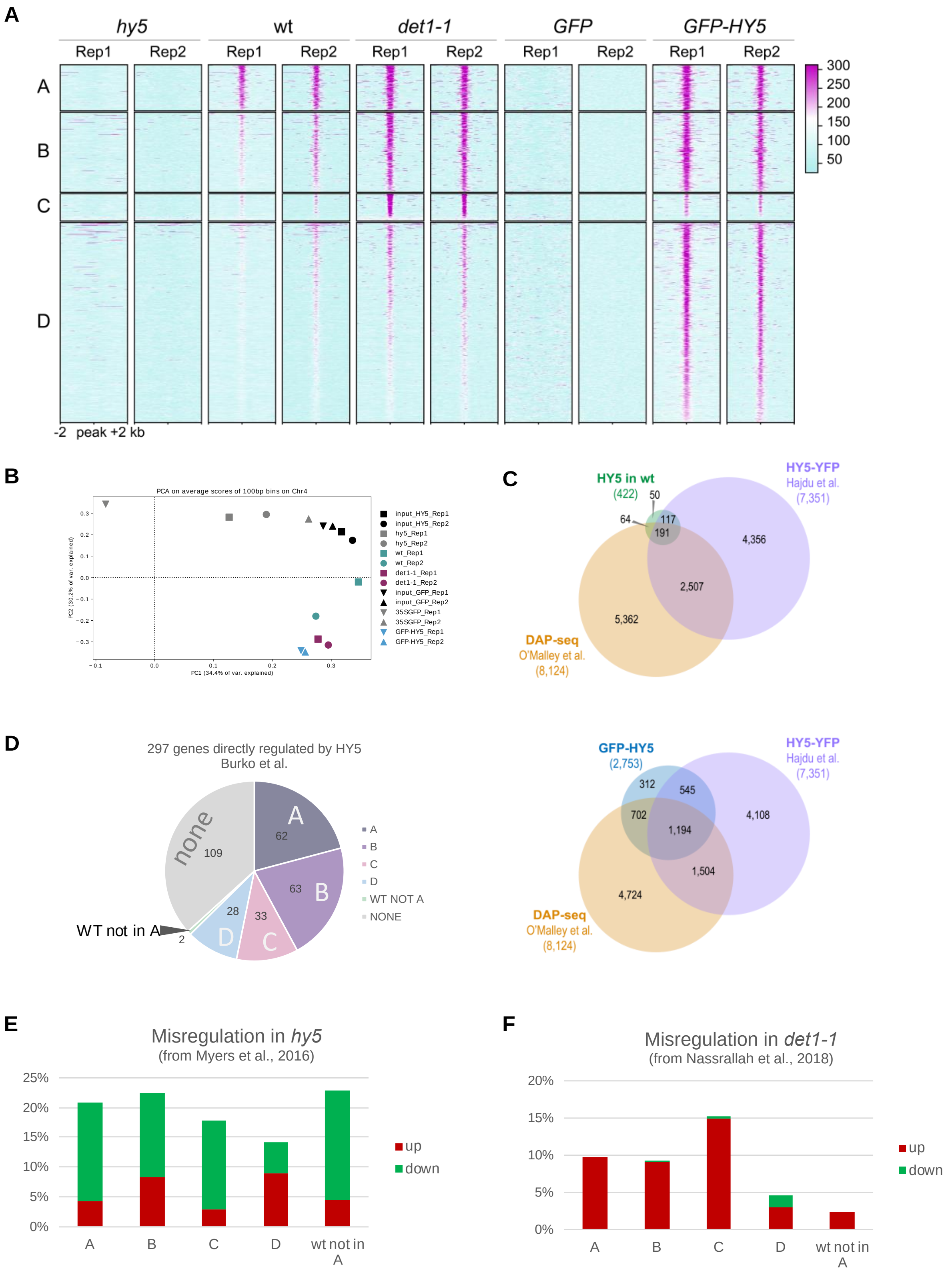

**Figure S3. Analysis of HY5 targets in comparison with previous published binding and expression data**
(A) Heatmaps showing the relative enrichment as in Fig. 5B with each biological replicate shown separately.
(B) Principal component analysis of the different samples used for the ChIP analysis.
(C) Comparisons of the HY5 endogenous and the overexpressed GFP-HY5 target genes with the HY5 targets found by ChIP-seq in *Hajdu et al*. (upper panel) and by DAP-seq in O'Malley *et al*. 2014 (lower panel).
(D) Proportion of genes belonging to the A, B, C and D classes among the genes found to be directly regulated by HY5 in Burko *et al*. 2020.
(E and F) Genes belonging to the A, B, C and D classes overlapping with genes misregulated in *hy5* in Myers *et al*., 2016 (E) or with genes misregulated in *det1-1* in Nassrallah *et al*., 2018 (F).

**Class A.** 412 peaks/378 genes

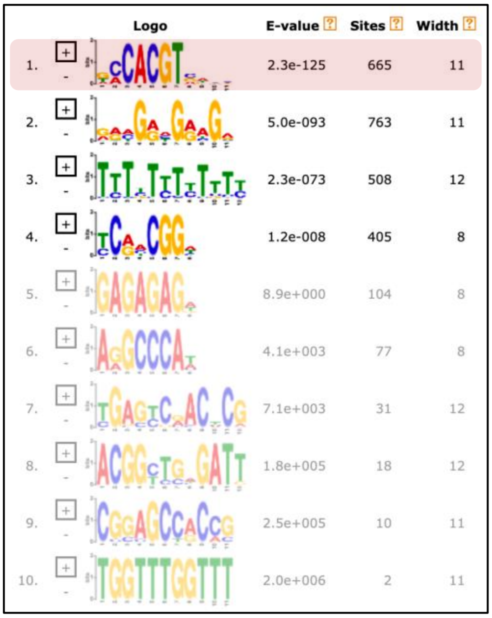

**Class B.** 715 peaks/649 genes

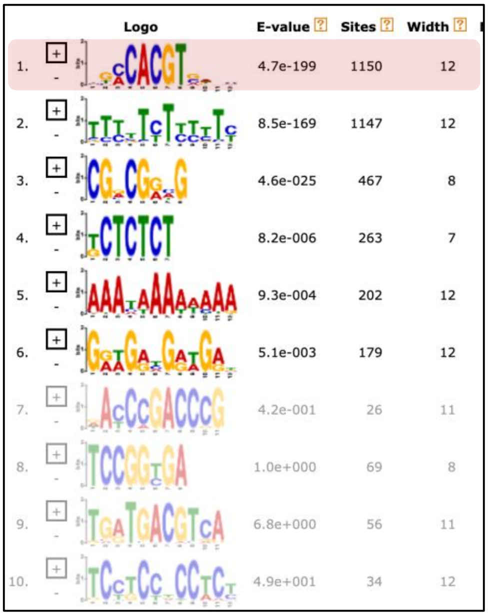

**Class C.** 245 peaks/236 genes

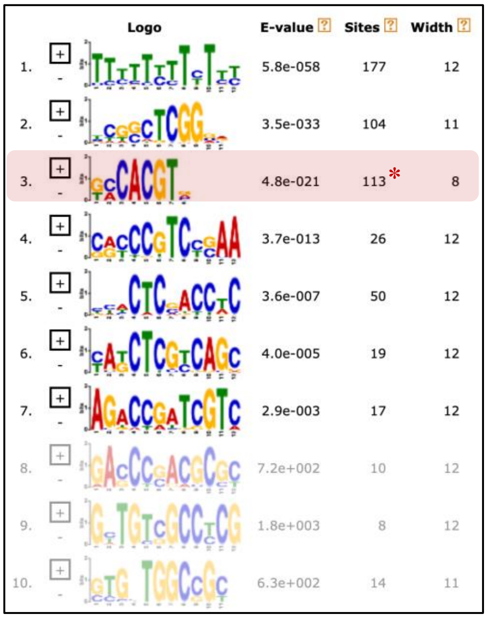

**Class D.** 1,767 peaks/1,720 genes

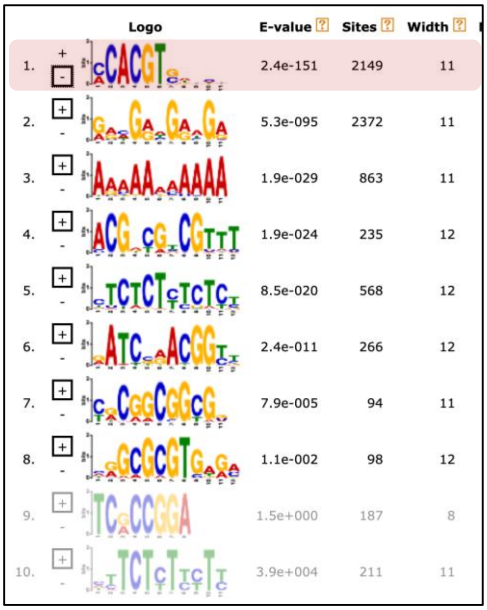

**Figure S4. *De novo* motif search under HY5 peaks annotated A to D gene classes.** The 10 motif found by the MEME software are shown for each class. A G-box (highlighted in red) was found as the top enriched motif with a number of occurrences of more than one motif per peak (number of sites > number of peaks) for all classes but C. Instead, for peaks on class C genes, the G-box motif was found with a high *E-value* under less than half of the peaks (*).

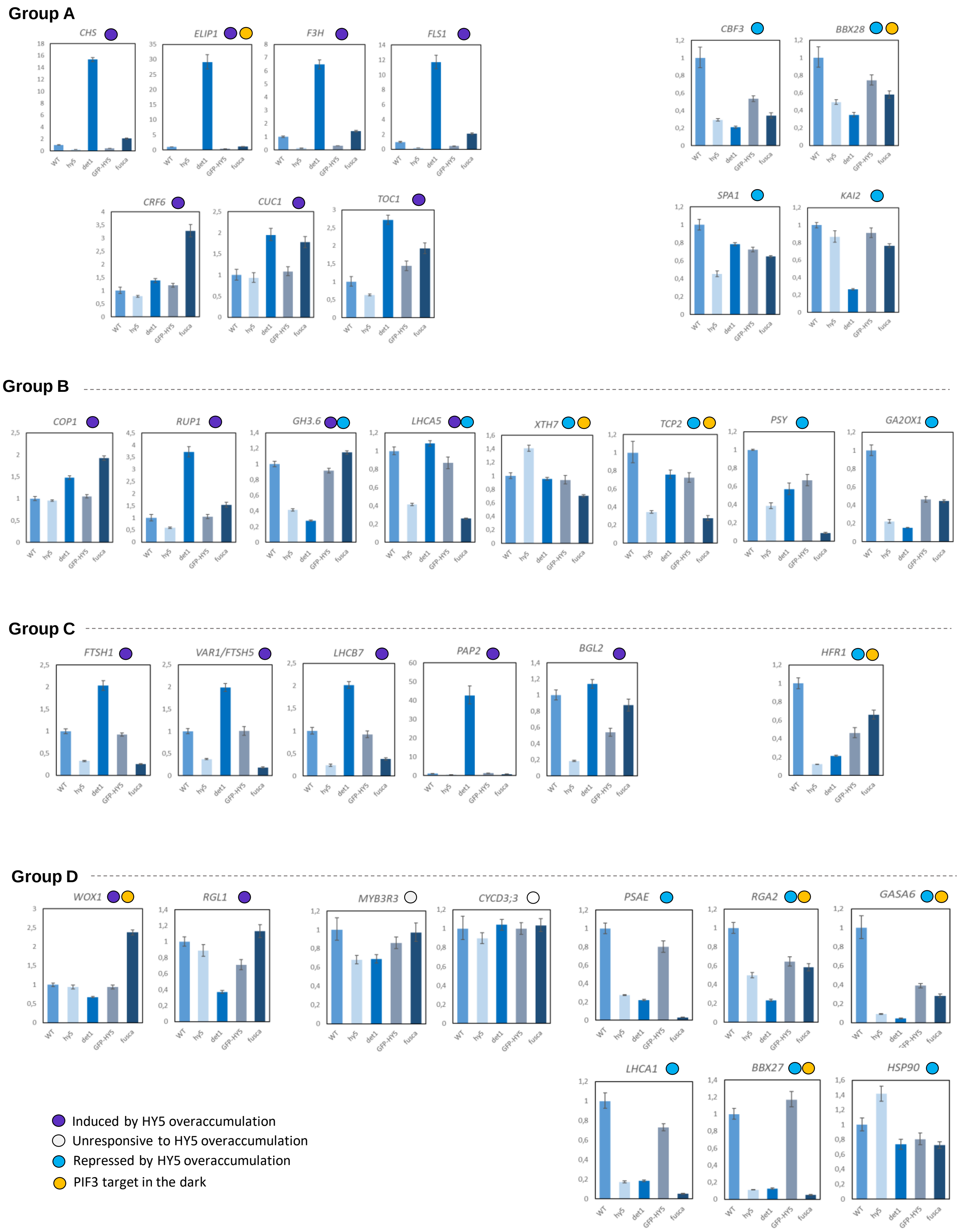

Group A
CHS
ELIP1
F3H
FLS1
CBF3
BBX28
CRF6
CUC1
TOC1
SPA1
KAI2
Group B
COP1
RUP1
GH3.6
LHCA5
XTH7
TCP2
PSY
GA2OX1
Group C
FTSH1
VAR1/FTSH5
LHCB7
PAP2
BGL2
HFR1
Group D
WOX1
RGL1
MYB3R3
CYCD3;3
PSAE
RGA2
GASA6
LHCA1
BBX27
HSP90
WT
hy5
det1
GFP-HY5
fusca
Induced by HY5 overaccumulation
Unresponsive to HY5 overaccumulation
Repressed by HY5 overaccumulation
PIF3 target in the dark

**Figure S5. Gene expression analysis of HY5 bound genes.**
Relative accumulation of transcripts in Arabidopsis seedlings grown in long day and collected in light, analysed by qRT-PCR. The expression level of each gene was normalized to that of *PP2A*. Expression levels for each gene are shown relative to the expression levels in WT, which is set as 1. Bars represent average ± SD obtained from three replicates and the experiment was repeated with different pools of plants with similar results. The genes are grouped by the classes defined in Fig. 5A; GFP-HY5 are complemented plants while *fusca* means the *fusca*-like plants from the same GFP-HY5/*hy5* transgenic line.

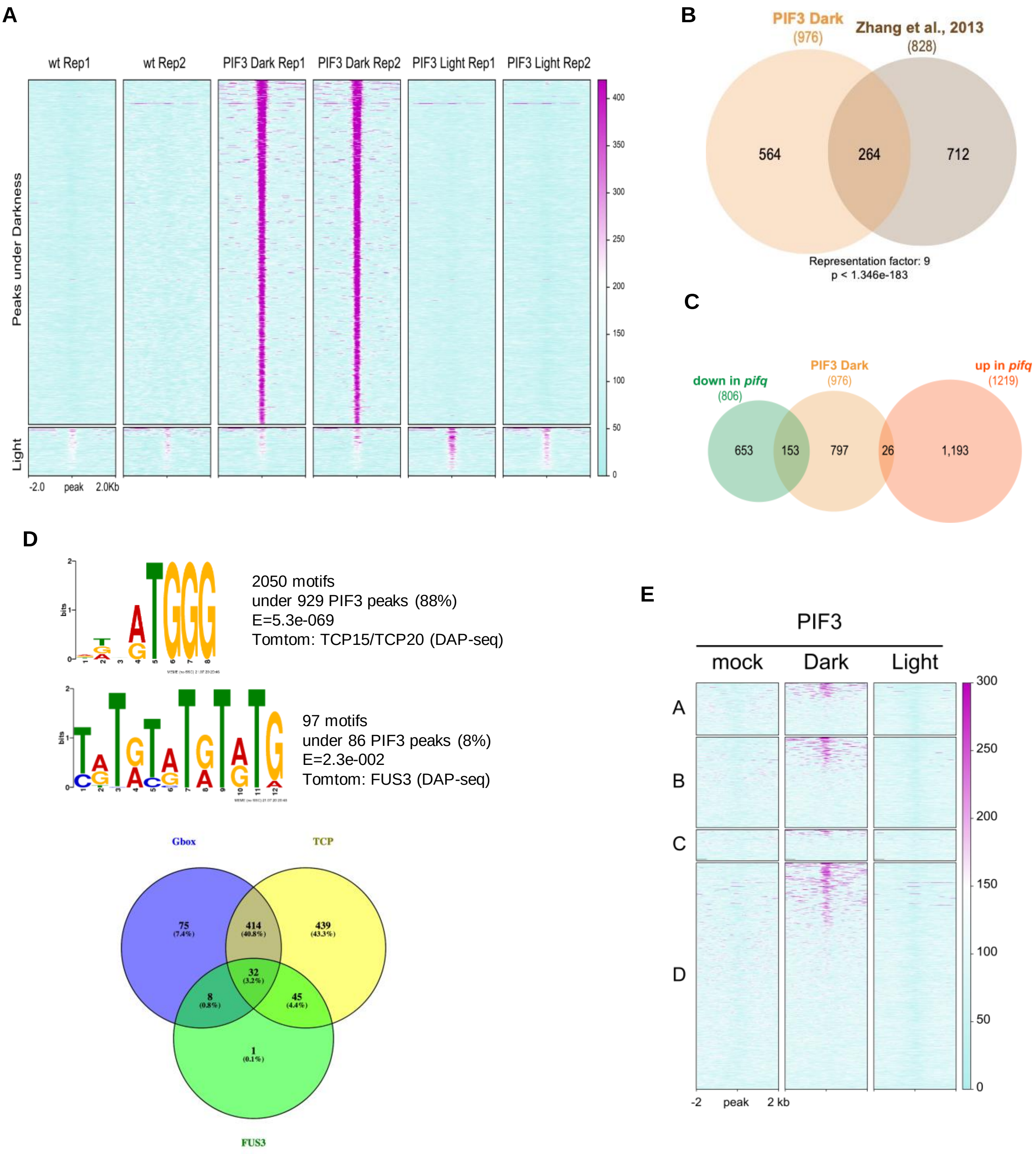

**Figure S6. Analysis of PIF3 targets expression and binding sites and overlapping with HY5 binding classes.**
(A) Heatmaps showing the relative enrichment as in Fig. 7A with each biological replicate shown separately.
(B) Venn diagram showing the overlap between the sets of PIF3-bound genes found in this study and in Zhang *et al*., 2013.
(C) Venn diagram showing the overlap between the sets of PIF3-bound genes found in this study and misregulated genes in *pifq* (Zhang *et al*., 2013).
(D) Extra motifs found under the PIF3 peaks in dark conditions.
(E) Heatmaps showing the relative enrichment of PIF3 around the HY5 peaks found in the promoters of the A, B, C and D gene classes. Genes in each class are ranked according to the PIF3 signal.

**Table S1. Interactomics of DET1 and COP1 proteins.**

List of proteins that associate with DET1 or COP1 in TAP assays performed in Arabidopsis cell cultures. Number of peptides for each experiment is depicted. Proteins are associated by function or functional complexes and a colour code is used to show proteins that associate with DET1 and COP1 (blue); that were found only in association with DET1 (green) or COP1 (orange). The maximum number of peptides represented were used as the base for **Fig.1** representation.

| | | | DET1 TAPs | | | | COP1 TAPs | | |
|---|---|---|---|---|---|---|---|---|---|
| | Protein | Gene ID | Appearance in nº experimets (max=5) | Sum of peptides (all exp.) | Max nº of peptides in one exp. | | Appearance in nº experimets (max=2) | Sum of peptides (all exp.) | Max nº of peptides in one exp. |
| CRL4$^{C3D}$ complex | DET1 | AT4G10180 | 5 | 318 | 103 | | 2 | 7 | 4 |
| | DDB1a | AT4G05420 | 5 | 403 | 134 | | 2 | 11 | 8 |
| | DDB1b | AT4G21100 | 5 | 413 | 152 | | 1 | 7 | 7 |
| | COP10 | AT3G13550 | 5 | 21 | 7 | | 0 | 0 | 0 |
| | DDA1 | AT5G41560 | 5 | 20 | 12 | | 0 | 0 | 0 |
| | CUL4 | AT5G46210 | 5 | 44 | 12 | | 0 | 0 | 0 |
| COP signalosome | CSN1 | AT3G61140 | 5 | 22 | 9 | | 0 | 0 | 0 |
| | CSN2 | AT2G26990 | 3 | 10 | 4 | | 0 | 0 | 0 |
| | CSN3 | AT5G14250 | 1 | 3 | 3 | | 0 | 0 | 0 |
| | CSN4 | AT5G42970 | 2 | 5 | 4 | | 0 | 0 | 0 |
| | CSN5A | AT1G22920 | 2 | 2 | 1 | | 0 | 0 | 0 |
| | CSN5B | AT1G71230 | 0 | 0 | 0 | | 0 | 0 | 0 |
| | CSN6A | AT5G56280 | 3 | 5 | 2 | | 0 | 0 | 0 |
| | CSN6B | AT4G26430 | 0 | 0 | 0 | | 0 | 0 | 0 |
| | CSN7 | AT1G02090 | 3 | 3 | 1 | | 0 | 0 | 0 |
| | CSN8 | AT4G14110 | 1 | 1 | 1 | | 0 | 0 | 0 |
| COP1-SPA complexes | COP1 | AT2G32950 | 3 | 6 | 4 | | 2 | 158 | 86 |
| | SPA1 | AT2G46340 | 2 | 14 | 11 | | 2 | 13 | 11 |
| | SPA2 | AT4G11110 | 0 | 0 | 0 | | 2 | 14 | 12 |
| | SPA3 | AT3G15354 | 0 | 0 | 0 | | 2 | 27 | 20 |
| | SPA4 | AT1G53090 | 0 | 0 | 0 | | 2 | 103 | 70 |
| Light percpt. | CRY1 | AT4G08920 | 0 | 0 | 0 | | 2 | 17 | 9 |
| | CRY2 | AT1G04400 | 3 | 7 | 4 | | 2 | 15 | 12 |

**Table S2. List of DET1 and COP1 associated proteins found in the different replicates of TAP assays.**
TAP assays were performed in Arabidopsis cell cultures grown first in the dark and then kept under dark conditions (DET1 TAP assays #1, #2, #3, #4 and COP1 TAP assay #2 cells) or transferred to light (DET1 TAP assay #4 and COP1 assay #1) for 24h before cells were harvested. Protein score and number of peptides for each experiment is depicted. Proteins are associated by function or functional complexes and a colour code is used to show proteins that associate with DET1 and COP1 (blue); that were found only in association with DET1 (green) or COP1 (orange).

| | | | DET1 TAP-tag | | | | | COP1 TAP-tag | |
|---|---|---|---|---|---|---|---|---|---|
| | Protein | Gene ID | PROTEIN SCORE/ Nº PEPTIDES | | | | | PROTEIN SCORE/ Nº PEPTIDES | |
| | | | Assay 1 | Assay 2 | Assay 3 | Assay 4 | Assay 5 | Assay 1 | Assay 2 |
| CRL4$^{C3D}$ complex | DET1 | AT4G10180 | 274/26 | 409/30 | 1189/91 | 2763/103 | 2035/68 | 136/4 | 92/3 |
| | DDB1a | AT4G05420 | 303/42 | 522/46 | 1913/134 | 3243/121 | 1624/60 | 275/8 | 106/3 |
| | DDB1b | AT4G21100 | 301/44 | 466/48 | 1931/103 | 4097/152 | 1697/66 | 247/7 | 0 |
| | COP10 | AT3G13550 | 14/4 | 19/5 | 198/7 | 64/2 | 63/3 | 0 | 0 |
| | DDA1 | AT5G41560 | 3/1 | 3/1 | 279/12 | 157/3 | 112/3 | 0 | 0 |
| | CUL4 | AT5G46210 | 27/11 | 26/12 | 303/7 | 240/8 | 147/6 | 0 | 0 |
| COP signalosome | CSN1 | AT3G61140 | 15/6 | 11/5 | 71/1 | 34/1 | 59/9 | 0 | 0 |
| | CSN2 | AT2G26990 | 10/4 | 10/4 | 71/2 | 0 | 0 | 0 | 0 |
| | CSN3 | AT5G14250 | 0 | 0 | 124/3 | 0 | 0 | 0 | 0 |
| | CSN4 | AT5G42970 | 0 | 0 | 159/4 | 48/1 | 0 | 0 | 0 |
| | CSN5A | AT1G22920 | 0 | 0 | 41/1 | 52/1 | 0 | 0 | 0 |
| | CSN5B | AT1G71230 | 0 | 0 | 0 | 0 | 0 | 0 | 0 |
| | CSN6A | AT5G56280 | 6/2 | 3/1 | 66/2 | 0 | 0 | 0 | 0 |
| | CSN6B | AT4G26430 | 0 | 0 | 0 | 0 | 0 | 0 | 0 |
| | CSN7 | AT1G02090 | 2/1 | 2/1 | 31/1 | 0 | 0 | 0 | 0 |
| | CSN8 | AT4G14110 | 0 | 0 | 27/1 | 0 | 0 | 0 | 0 |
| COP1-SPA complexes | COP1 | AT2G32950 | 2/1 | 8/4 | 38/1 | 0 | 0 | 2325/86 | 1749/72 |
| | SPA1 | AT2G46340 | 6/3 | 23/11 | 0 | 0 | 0 | 597/11 | 115/2 |
| | SPA2 | AT4G11110 | 0 | 0 | 0 | 0 | 0 | 526/12 | 73/2 |
| | SPA3 | AT3G15354 | 0 | 0 | 0 | 0 | 0 | 196/7 | 654/20 |
| | SPA4 | AT1G53090 | 0 | 0 | 0 | 0 | 0 | 1116/33 | 1653/70 |
| Light perct. | CRY1 | AT4G08920 | 0 | 0 | 0 | 0 | 0 | 359/8 | 392/9 |
| | CRY2 | AT1G04400 | 0/1 | 7/4 | 73/2 | 0 | 0 | 463/12 | 68/3 |

**Table S3. List of DWD proteins that associate with DET1.**

Protein score and number of peptides are displayed. These DWD proteins were listed in Lee *et al.*, 2007 and reviewed in Fonseca and Rubio, 2019.

| TAP-tag DET1 | | Assay 1 | Assay 2 | Assay 3 | Assay 4 | Assay 5 |
|---|---|---|---|---|---|---|
| **Protein** | **Gene ID** | | | | | |
| PWP2 | AT1G15440 | 0 | 0 | 0 | 34/1 | 0 |
| SCD1 | AT1G49040 | 2,3/2 | 7,42/3 | 0 | 0 | 0 |
| protein with a DWD motif | AT1G65030 | 1,76/2 | 0/1 | 0 | 0 | 0 |
| SMU1 | AT1G73720 | 2,04/1 | 0 | 0 | 0 | 0 |
| Transducin/WD40 | AT1G78070 | 0 | 0 | 21,0/1 | 0 | 0 |
| MSI4 | AT2G19520 | 0 | 0/2 | 0 | 0 | 0 |
| MAC3B | AT2G33340 | 4,73/2 | 7,32/3 | 0 | 0 | 0 |
| Transducin/WD40 | AT2G43770 | 0 | 0 | 0 | 0 | 31,0/1 |
| TIF3I1 | AT2G46280 | 0 | 0 | 62/1 | 0 | 0 |
| Transducin/WD40 | AT2G46560 | 0 | 2,62/1 | 0 | 0 | 0 |
| SWA1, EDA13, EDA19 | AT2G47990 | 0/1 | 1,80/1 | 0 | 0 | 0 |
| RAPTOR1 | AT3G08850 | 0 | 0/1 | 0 | 0 | 0 |
| BUB3 | AT3G19590 | 1,85/1 | 0 | 0 | 0 | 0 |
| RID3 | AT3G49180 | 0 | 2,32/1 | 0 | 0 | 0 |
| VIP3 | AT4G29830 | 2,07/1 | 0 | 0 | 0 | 0 |
| FY | AT5G13480 | 0 | 2,30/1 | 0 | 0 | 0 |
| APRF1; S2LA | AT5G14530 | 2,2/1 | 0 | 0 | 0 | 0 |
| PEP2 | AT5G15550 | 0 | 2,16/1 | 0 | 0 | 0 |
| MSI1 | AT5G58230 | 2,57/1 | 2,96/1 | 0 | 0 | 0 |
| PRL2 | AT3G16650 | 0 | 0 | 20,0/1 | 0 | 0 |

**Table S4. Transcription factors that are direct targets of HY5 in wild-type.**

| Family | Locus | Name | Description |
|---|---|---|---|
| **AP2/EREBP** | AT4G25480 | CBF3 | This gene is involved in response to low temperature and abscisic acid |
| | AT1G75490 | - | This gene is involved in response to drought |
| | AT1G01250 | - | - |
| | AT5G47220 | ERF2 | ETHYLENE RESPONSE FACTOR- 2 |
| | AT1G51190 | PLT2 | PLETHORA 2 |
| | AT3G61630 | CRF6 | CYTOKININ RESPONSE FACTOR 6 |
| | AT5G60120 | TOE2 | TARGET OF EARLY ACTIVATION TAGGED (EAT) 2 |
| **bZIP** | AT4G34000 | ABF3 | ABSCISIC ACID RESPONSIVE ELEMENTS-BINDING FACTOR 3 |
| | AT3G19290 | ABF4 | ABA-RESPONSIVE ELEMENT BINDING PROTEIN 2 |
| | AT4G01120 | GBF2 | G-BOX BINDING FACTOR 2 |
| | AT2G46270 | GBF3 | G-BOX BINDING FACTOR 3 |
| | AT5G11260 | HY5 | ELONGATED HYPOCOTYL 5 |
| | AT3G17609 | HYH | HY5-HOMOLOG |
| | AT1G45249 | ABF2 | ABSCISIC ACID RESPONSIVE ELEMENTS-BINDING FACTOR 2 |
| | AT1G19490 | - | Basic-leucine zipper (bZIP) transcription factor |
| **NAC** | AT3G15170 | CUC1 | CUP-SHAPED COTYLEDON1 |
| | AT4G01540 | NTM1 | NAC WITH TRANSMEMBRANE MOTIF1 regulates cell division in Arabidopsis |
| | AT5G18270 | ANAC087 | NAC domain containing protein 87 |
| **bHLH** | AT1G05805 | AKS2 | ABA-RESPONSIVE KINASE SUBSTRATE 2 |
| | AT1G62975 | - | basic helix-loop-helix (bHLH) DNA-binding |
| | AT1G59640 | BPE | BIGPETAL is involved in the control of petal size |
| **MYB** | AT2G47460 | MYB12 | PRODUCTION OF FLAVONOL GLYCOSIDES 1 (PFG1) |
| | AT1G35515 | MYB8 | High response to osmotic stress 10 |
| **MYB-related** | AT4G01280 | RVE5 | REVEILLE 5 plays a minor role in clock regulation. |
| | AT1G01520 | RVE3 | ALTERED SEED GERMINATION 4 |
| | AT1G17460 | TRFL3 | TRF-LIKE 3 |
| | AT5G58900 | DIV1 | R-R-type MYB protein |
| **G2-like** | AT3G46640 | LUX | LUX ARRHYTHMO |
| | AT4G18020 | PRR2 | PSEUDO-RESPONSE REGULATOR 2 |
| **C2C2-Dof** | AT5G60850 | OBP4 | OBF BINDING PROTEIN 4 |
| **C2C2-CO-like** | AT4G27310 | BBX28 | Negative regulator of photomorphogenic development |
| **C3H** | AT2G41900 | OXS2 | OXIDATIVE STRESS 2 |
| **SRS** | AT5G66350 | SHI | SHORT INTERNODES is involved in the response to gibberellic acid |
| | AT3G51060 | STY1 | STYLISH 1 |
| **SBP** | AT1G02065 | SPL8 | SQUAMOSA PROMOTER BINDING PROTEIN-LIKE 8 |
| **Trihelix** | AT1G76890 | GT2 | Encodes a plant trihelix DNA-binding protein |
| **CAMTA** | AT4G16150 | CAMTA5 | CALMODULIN-BINDING TRANSCRIPTION ACTIVATOR 2 |
| **HFS** | AT3G02990 | HSFA1E | HEAT SHOCK TRANSCRIPTION FACTOR A1E |
| | AT5G16820 | ATHSF3 | ARABIDOPSIS HEAT SHOCK FACTOR 3 |
| **TAZ** | AT4G37610 | BT5 | BTB AND TAZ DOMAIN PROTEIN 5 |
| **PHD** | AT1G77800 | PRE2 | PACLOBUTRAZOL RESISTANCE2 |
| **E2F-DP** | AT1G47870 | E2FC | ARABIDOPSIS THALIANA HOMOLOG OF E2F C |
| **CPP** | AT4G14770 | TCX2 | TESMIN/TSO1-LIKE CXC |
| **Pseudo ARR-B** | AT5G61380 | TOC1 | TIMING OF CAB EXPRESSION 1 |
| | AT2G46790 | TL1 | TOC1-LIKE PROTEIN 1 |
| **NF-Y** | AT5G47640 | NFY-B2 | NUCLEARFACTOR Y, SUBUNIT B2 |

### Table S5. Oligonucleotides used in this study

| Forward | SEQUENCE | Reverse | SEQUENCE |
|---|---|---|---|
| **SPA1 fw** | AGCTTTGGAGCATCAATGAGA | **SPA1 rv** | ATTGCACGCAACACACATTT |
| **TCP2 fw** | CGTCACCTACTACTACTAACTCCAAGC | **TCP2 rv** | GAAATGATTTTTAACCACAAGCAG |
| **CHS fw** | GAGAAGTTCAAGCGCATGTG | **CHS rv** | CGCTTCTTTGCCTAGCTTAG |
| **FTSH1 fw** | GAAGAAGAATGCAGTTGTCTCTGA | **FTSH1 rv** | CACTAGAGCATGACCAGCCTC |
| **F3H fw** | GTCTCTAGTCACCTCCAGG | **F3H rv** | AGCCAAACTCATAAGCCTC |
| **PAP2 fw** | GAGGAAAGGTGCATGGACTG | **PAP2 rv** | TCTTTCTGCATCGATTTAGCC |
| **ELIP1fw** | TTCACCAGCACCATCTACCTC | **ELIP1 rv** | CGCTAAACTTTGTGCTCACCTT |
| **HFR1 fw** | TTCAGTTACTCGAAAAGGTTCCA | **HFR1 rv** | CGAAACCTTGTCCGTCTTG |
| **FLS1 fw** | CATCGGCGATCAGATTCTGAG | **FLS1 rv** | GGGAGGCTCCAAGAAAACC |
| **VAR1 fw** | GATTGGACAAGTTGCAGTTGG | **VAR1 rv** | GAGTAATCCTTCTGCGATGACATA |
| **KAI2 fw** | GATCTCTGCTTCTCCGAGATA | **KAI2 rv** | GAGCGAAACCTAAGCACCAC |
| **BGL2 fw** | GAGGTGGAGACACCACGAGT | **BGL2 rv** | TCTGCGGCGTGTGTAGTC |
| **BBX28 fw** | TCCTCCGGTGATGAGTTCTT | **QBBX28 rv** | GCTGAAGAGCTTCCGATCTC |
| **LHCB7 fw** | TCCAAGCTTCAGGGAGAGAC | **LHCB7 rv** | CTTGCGTATTCCGGTCCTAC |
| **CUC1 fw** | AGTCTGAGCCTTGGGAGCTT | **CUC1 rv** | AGCTTCTGTTGCTCTGTTCGT |
| **LHCA1 fw** | CCGGGAATGTTGGTCGTAT | **LHCA1 rv** | GGTCAAACCCAAAGTCACCA |
| **TOC1 fw** | CGAGCTTCTGAACTTGTGGAC | **TOC1 rv** | TTCTTCTCAGCAAGTCCTAGCAT |
| **WOX1 fw** | CCACGGATTCGACAAGAAA | **WOX1 rv** | GATGATAATCACGGGAAGGTTT |
| **CBF3 fw** | CGCTAAGGACATCCAAAAGG | **CBF3 rv** | CGCATCACACATCTCATCCT |
| **GASA6 fw** | AGAAACCCCAATCTGTTTCC | **GASA6 rv** | GAAGGTCCATACACATTTCG |
| **CRF6 fw** | TCGAGCCGTACTCGGATTT | **CRF6 rv** | TCGTACGCTATAGCAGCTTCC |
| **BBX27 fw** | CGAATGGTTAAACCTAAGGTGC | **BBX27 rv** | CACCTCGTTCTGTCAAGGAAG |
| **PSY fw** | TCAGAGACGTAGGCGAAGA | **PSY rv** | TCTCAGCTTCGTCGAAGAA |
| **RGA2 fw** | AACTCGGCATGTTGTCCTG | **RGA2 rv** | AAAGCGCGTGAACGAGAC |
| **LHCA5 fw** | GATTCGTAATTTACCGGTTTGG | **LHCA5 rv** | CCTTTTAGTCTCAGCAAACCCCAT |
| **RGL1 fw** | CAAGCATGTTGTTGGCACTT | **RGL1 rv** | GCAACAAACAACCTTCATTCTCT |
| **GH3.6 fw** | CATCTCTGAGTTCCTCACAAGTTC | **GH3.6 rv** | CCAGGAACAAACTGGTCCAT |
| **CYCD3,3 fw** | GTTCCTTTTCTCTTAGATTTTCAAGTG | **CYCD3,3 rv** | AACGAGATTGGAGTCACAGGA |
| **COP1 fw** | TGGTAGTGACGACTGCAAGGTT | **COP1 rv** | GAGCCAGGATTGTACTTGACACA |
| **PSAE fw** | CGGCTTCCTCTTCTTCAAAA | **PSAE rv** | GGATTCTCTCCTTAGAATCTTGACCT |
| **XTH7 fw** | CAAATTGGACCCTAGCTCAGGAT | **XTH7 rv** | GCAGTGACAGTCCGGCAGAA |
| **HSP90 fw** | CGAGTTTGGCAAGTCTGTCA | **HSP90 rv** | TTTCCATCGGACTTTGTTGTC |
| **GA2OX1 fw** | CTCCCGATCACACTTCCTTC | **GA2OX1 rv** | CCTCCCATTTGTCATCACCT |
| **MYB3R3 fw** | CTCCGGAAGAGGATGAGACA | **MYB3R3 rv** | CGATGCAGGCATTGAACTT |
| **RUP1 fw** | ACGTAACGGAGGGACGTTAG | **RUP1 rv** | CGGATCGAACTCTACCGAAC |
| **PP2A fw** | TAACGTGGCCAAAATGATGC | **PP2A rv** | GTTCTCCACAACCGCTTGGT |
| **HY5 fw** | AGAACAAGCGGCTGAAGAGGT | **HY5 rv** | TGTCTAAGCATCTGGTTCTCGT |
| **ACT8 fw** | GACTCAGATCATGTTTGAGACCTTT | **ACT8 rv** | TCACCAGAGTCCAACACAATACC |
